# Anti-NMDAR encephalitis antibodies cause long-lasting degradation of the hippocampal neural representation of memory

**DOI:** 10.1101/2022.11.25.517901

**Authors:** AmirPasha Zamani, Paula Peixoto-Moledo, David P. Tomàs, Horacio G. Rotstein, Josep Dalmau, Pablo E. Jercog

**Affiliations:** Institut d’Investigacions Biomèdiques August Pi i Sunyer (IDIBAPS), Barcelona, Spain, 08036; Federated Department of Biological Sciences, New Jersey Institute of Technology and Rutgers University, Newark, NJ 07102, U.S.A.; Department of Neurology, Perelman School of Medicine, University of Pennsylvania, Philadelphia, US, 19104; Institució Catalana de Recerca i Estudis Avançats (ICREA), Barcelona, Spain, 08010; Neurology Department, Institute of Neuroscience, Hospital Clínic de Barcelona, Barcelona, Spain 08036

**Keywords:** anti-NMDAR-encephalitis, calcium-imaging, hyperactivity, amnesia, mechanistic

## Abstract

N-methyl D-aspartate receptor (NMDAR) encephalitis is an immune-mediated disorder characterized by a complex neuropsychiatric syndrome together with a reduction of NMDAR. Although in most patients the life-threatening symptoms of the acute stage resolve with immunotherapy, memory and executive functions remain altered for several months or years. A mechanistic explanation for these long-lasting cognitive effects is still lacking and previous animal models have not explored this effect.

Here, we combined repeat calcium imaging of the same population of hundreds of hippocampal CA1 neurons for three months along with two behavioral tasks to assess retrograde and anterograde memory loss using a reported mouse model of cerebroventricular transfer of patients’ CSF antibodies. We measured how memory-related neuronal activity is affected by the presence of NMDAR antibodies during the induction of the model and its long-lasting recovery. In addition, we developed a computational model that provides a mechanistic explanation for the long-term antibody-mediated impairment of memory.

The findings show that the presence of antibodies leads to an increase of CA1 neuronal firing rate, resulting in a reduction of the amount of information encoded by these cells. Furthermore, the antibodies cause a degradation of the hippocampal neuronal response stability over time, providing a neural correlate of memory dysfunction. All these neuronal alterations span the 3 months of recordings, and in some cases beyond the last recording point. The computational model shows that a reduction of NMDAR is sufficient to cause the changes observed in neuronal activity, including the different involvement of excitatory and inhibitory inputs to CA1 neurons.

Altogether, we show that the antibody-mediated reduction of NMDAR leads to long-term changes in hippocampal neuronal activity which extend far beyond the antibody clearance, providing a mechanism that can account for the cognitive deficits observed in the protracted recovery of patients with anti-NMDAR encephalitis.

## Introduction

N-methyl D-aspartate receptor (NMDAR) encephalitis (anti-NMDARe) is the most common neuronal antibody-mediated encephalitis.^1^ It is associated with antibodies against the GluN1 subunit of the NMDAR. The disorder is characterized by a complex neuropsychiatric syndrome that in the acute phase associates with prominent behavioral and cognitive dysfunction along with short-term memory loss, decreased level of consciousness, seizures, abnormal movements, dysautonomia, and central hypoventilation that often requires intensive care.^2^ This phase is followed by a prolonged post-acute stage characterized by the resolution of many of the severe neurologic symptoms, while cognitive functions (particularly verbal and visuospatial memory) and executive function tend to improve, but a much slower pace, resulting in substantial disability for several months or even years.^3-7^ The mechanisms underlying the long-lasting cognitive deficits are unclear.

In-vitro studies have shown that the interaction between the antibodies and NMDAR leads to a reduction of synaptic NMDARs and NMDA-mediated currents in the hippocampus.^8^ In vivo studies using a mouse model of continuous infusion of CSF from patients with anti-NMDARe showed that this decrease in the density of total and synaptic NMDAR was correlated with progressive memory deficits and other behavioral changes that resemble different features of the anti-NMDARe in patients.^9^ However, in this mouse model, the behavioral alterations and tissue reduction of synaptic NMDAR resolve soon after the infusion is stopped.

It remains unclear whether the long-term memory impairment observed in patients is due to the presence of antibody-producing cells, or more generally, mediators of inflammation. The alterations of memory caused by the reduction of NMDAR and their neuronal correlates have been extensively studied in non-immunological paradigms.^10,11^ It has been previously suggested that the reduction of NMDAR can drastically change the neuronal network activity.^12^ However, it is not known if a rapid change in NMDAR density during a short time window (i.e., 2 weeks as in the model of^9^) can lead to long-term changes at the network level. We hypothesize that a short hippocampal exposure to patients’ NMDAR antibodies leads to a long-lasting disruption of the network state, not only affecting the formation of new memories but also altering the ability to recall previously stored memories. A better understanding of the mechanisms underlying memory dysfunction during anti-NMDARe may provide new treatment strategies for the memory and behavioral alterations of patients with the disease.

Traditional electrophysiology is not suitable for assessing the learning of long-term memories mediated by NMDAR,^13^ a process that involves large populations of neurons and lasts from hours to days.^14^ In contrast, calcium imaging allows recording the same population of neurons over periods that span days or weeks.^15^ Moreover, miniaturized microscopes used to register hippocampal calcium imaging allow neuronal activity recordings while the animal freely moves to solve spatial memory tasks, which is an ethological form of memory in rodents. This combination of techniques offers a unique opportunity to study memory in animal models over the time scale of memory formation. For this study, we collected unprecedented data sets of hundreds of neurons per animal for approximately 3 months, to study the acute and long-lasting mnestic dysfunction along with the underlying alterations in hippocampal neuronal function in animals treated with patients NMDAR antibodies and appropriate controls.

## Materials and methods

### Ethical Aspects

The study was approved by the local institutional review board (Hospital Clínic, HCB/2018/0192). Informed consent was obtained from all patients for the use of CSF for research purposes. All animal procedures were approved and executed following institutional guidelines (Generalitat de Catalunya: Autorització de projecte d’experimentació N? = 9997), and the Comitè Ètic d’experimentació Animal (CEEA) from University of Barcelona, Nº = 121/18.

### Viral injection for calcium imaging

Virus injection surgeries were performed on male C57BL/6J mice (Janvier Labs) at 6-8 weeks of age. Animal housing and husbandry was reported in^9^. We labeled hippocampal CA1 excitatory neurons by injecting an adeno-associated virus (AAV, serotype 2.5), driving the expression of GCaMP6m via the CaMKIIα promoter, which limits expression to pyramidal neurons. The virus was injected via a borosilicate glass pipette with a 50 μm diameter tip using short pressure pulses applied with a picospritzer (Parker). Six hundred nanoliters were distributed equally in three different hippocampal locations at coordinates relative to bregma on the right hemisphere: [mediolateral (ML) = 1.8mm, anterior-posterior (AP) = −1.5, dorsoventral (DV) = −1.6; ML = 1.4., AP = −2.2, DV = −1.55; ML = 2.1., AP = −2.9, DV = 1.8, from bregma].

### Mini-microscope placement and intracranial catheters implantation

A month after viral injection, a mini-endoscope is implanted to generate a cylindrical window in direct contact with the brain tissue. The mini-endoscope allows the insertion of a GRIN lens (Grintech, 1mm diameter, 4mm length, n.a. 0.5) to directly observe the neurons under the surface of the mini-endoscope (see Suppl. Material). This lens is used to focus the mini-microscope used for calcium recordings. The mini-endoscope was then inserted at the position and angle that covered most of the flat area of the dorsal part of the hippocampal CA1 region centered within the three viral injection locations [relative to bregma ML = 2.1(+ 77º on the coronal plane), AP = −2.2, DV = −1.1(from dura)]. During the same surgery, bilateral catheters (#3260PDG-1.5/SPC, Gauge 26; length 3.5mm, Plastics One, Roanoke, VA) for intraventricular infusion of patients antibodies, were implanted into the lateral ventricles (Suppl. Fig. 1). A detailed description of this procedure is included in the Supplementary Material.

A month after catheter implantation and approximately 60 days after viral injection, a daily inspection of the virus expression was performed to determine if calcium imaging activity was sufficiently high and stable across days to perform optical recordings. Once the optimal conditions are reached (i.e. the use of only 10% of the stimulation light on the sample to observe calcium events, and being able to track the same neurons across sessions), we mount, using non-invasive surgery, baseplates for mini-microscope attachment. The baseplate allows the fast attachment of the microscope to the animal for each session, without the performance of a surgery or the use of anesthesia, keeping the focal point across days and allowing the recording of the same neurons over the entire experiment length of 3 months.

### Recordings of individual neuronal activity

A nVista miniaturized one-photon microscope^16^ was used to record neuronal activity via calcium imaging in freely moving mice (Fig. 1A). We collected neuronal activity square images of 0.5 × 0.5 mm of CA1 tissue (Fig. 1B, left) at 20 Hz using the calcium indicator GCaMP6m^17^ from the CA1 area of the hippocampus. Neuronal electrical activity transduced into changes in the light refraction of the calcium indicator allows the measurement of green light as a readout of calcium spikes activation in the perisomatic and somatic regions of transfected pyramidal neurons. The images of the emitted light are captured via a CMOS camera in the microscope that sends digital information of each captured frame through an acquisition board to a computer (Suppl. Fig. 3A). Movies of neuronal activity corresponding to 10-20 minutes of behavioral tests are stored and then processed off-line (Suppl. Fig. 3B-D). From the captured frames, single calcium events are obtained using state-of-the-art methods. First, individual frames are realigned using the NoRMCorre algorithm^18^ to compensate for the field of view movement raised from the relative position between the lens and the brain tissue that continuously changes due to brain motion during animal behavior (Suppl. Fig. 3C). Once frames are aligned, background subtraction is performed to generate frames where only the relative change in fluorescence is observed, ΔF(t)/F_0_, where F_0_ is the mean luminosity obtained by averaging the entire movie on a given pixel^19^ (see Suppl. Fig. 3C, right). Spatial masks corresponding to individual cells were identified by CellMax^20^, an established cell sorting algorithm that finds candidates for potential neurons using a model of photon emission allowing the disentangling when neurons have overlapping contours (Fig. 1B, left, Suppl. Fig. 3C, center-left). Once high-scored cells are sorted based on five different criteria including distance from neighboring cells, size, shape, rising and decay dynamics of calcium events’ shapes, and signal-to-noise ratio. The discrimination is done mostly automatically by our CARPA algorithm,^21^ but approximately ten to twenty percent is decided manually based on feature discrimination. Masks of individual cells are stored for different types of analyses (Suppl. Fig. 3B, filtered neurons (green)). Alignment across sessions using NoRMCorre was critical to follow the same neurons across experimental days (Suppl. Fig. 3B, non-linear matching). Once individual neurons’ calcium traces were obtained as the average light intensity within each cell mask, calcium spikes events were detected using the MLSpike algorithm^22^ (Fig. 1B, right, Suppl. Fig. 3E). The event rate (i.e. firing rate) was quantified by the number of detected calcium events per second, which has almost a one to one correspondence with the number of action potentials when the number of events is small.^17^ Finally, a percentage of outlier neurons were excluded based on signal quality (criteria explained in Supplementary Methods).

**Figure 1.**
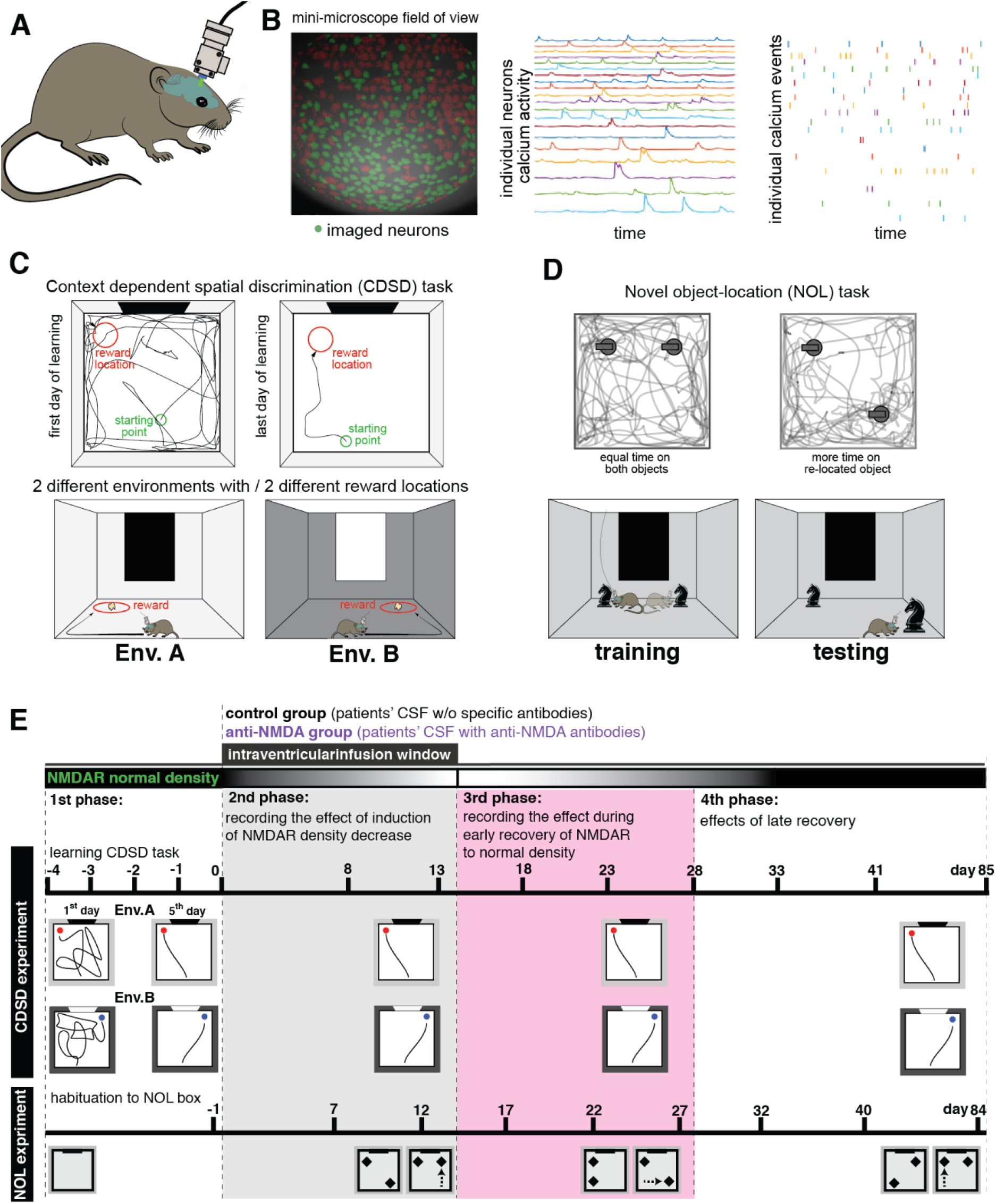
A novel experimental paradigm to observe the long-term effects of the immunoablation of NMDAR. **(A)** Schematic of the mouse with the head-mounted miniaturized one-photon microscope. A head-mounted mini microscope collects images of calcium concentration of the genetically labeled CA1 pyramidal cells while the animal can freely move to perform behavioral tasks. **(B) Left:** field of view of the calcium imaging miniaturized microscopy after the pre-processing. Our data analysis platform CaRPA provides the mask of the candidates of good-quality (green spots) or noisy neurons (red spots). **Center:** example traces of the selected good-quality neurons by our algorithm (see Methods). **Right:** calcium events (i.e. spikes) detected by our algorithm from the traces on the center panel(see Methods). **(C) Top Left:** mouse movement trajectories (gray) during the first session of learning the CDSD task (day-4). **Top Right:** mouse trajectory once the animal learned the CDSD task (day 0). **Bottom:** cartoon showing experimental boxes of the CDSD task, namely environments A and B. Boxes had different wall colors and visual cues. Food pellets were located at different relative locations, i.e., the northwest of env. A, and northeast of env.B. Mice were placed in the boxes at the central south point. **(D) Top Left:** example trace of the training session when the animals uniformly cover the arena and the two objects. **Top Right:** example trace of the testing session when the animal interacts more frequently with the object that has been relocated. **Bottom:** cartoon of the experiment where one can see the box and the two equal objects in the training session and then one is relocated in the testing session. **(E)** Schedule for behavior and neuronal activity recordings. In the first phase, all animals underwent a 5-day learning procedure for the behavioral tasks (see Methods). The 2nd phase is when the main experiment starts, infusing patients’ CSF and collecting the behavioral and neuronal activity data. The recovery period started on day 15 when the disease induction was stopped. Recovery consisted of two phases: the early recovery (first ten days shown in pink, the 3rd phase), and the late recovery (2 months, the 4th phase).

### Assessment of retro- and anterograde amnesia

#### Context-dependent spatial discrimination (CDSD) task to quantify the strength of old memories

We developed this task to quantify long-term spatial memories (Movie 1 (left panel day -4 right panel day 0)). The task is carried out in two 50 by 50 cm top-open boxes labeled Environment A and Environment B, respectively (Fig. 1C). Environment A has white walls, a textured floor surface, and a salty food pellet located on its northeast corner used as a reward. Environment B has black walls, a smooth floor surface, and a sweet food pellet on its northwest corner used as a reward (Fig. 1C, bottom). The same type of food corresponding to each box was grounded and uniformly spread over the entire box floor surface. Boxes were made of acrylic to help in cleaning and therefore avoid the odors of other mice that previously used the box. Both boxes have a cue card on the most distant wall from the experimenter (north wall), to provide orientation cues to the animal. Behavioral sessions lasted 10 minutes and started with Environment A immediately followed with Environment B. Only one session for each box was performed on each testing day.

Prior to each day of testing, mice were food-deprived for 16 hours to enhance their motivation for foraging. Animals were introduced near the center of the south wall and the time to reach the food pellet was used to quantify the animals’ spatial memory strength. Animals spatially navigated to maximize the harvesting of the food spread within the boxes in the 10 min sessions. Because of the size of the food pellet, mice develop a preference to find first its location (different in each of the boxes) and then continue foraging to eat the ground food that was spread throughout the box’s floor. Mice developed over days a stereotypical behavior characterized by shuttling directly from the starting location towards the food pellet location at the beginning of the 10 min sessions (Fig. 1C, top). Each environment (i.e. A and B) evoked a different shuttle trajectory (Fig. 1C, bottom). Formed memories were manifested when the shuttles became ballistic trajectories during the first few seconds of each session. Animals learned and displayed a ballistic trajectory over the 5 days of pre-training (i.e. days -4 to day 0). Utilizing the latency to find the regarded location, the CDSD task assesses the strength of the spatial memory learned by day 0, which is when the passive transfer of antibodies starts for the animal model’s induction. In this study, we call these CDSD memories: old memories.

#### Novel object location (NOL) task to quantify new memories

The NOL is a well-known task that measures animals’ newly formed spatially dependent memories (^23^, Fig. 1D, Movie 2). Briefly, the task consists of a 9-minute training session, and 4 hours later a 9-minute testing session. A discrimination index (DI) to measure memory performance, is defined as the difference between the exploration time of an old and a newly located object, divided by the total amount of time spent on exploring both objects in the arena. Animals were familiarized with the boxes (40 by 40 cm top-open) during the days before day 0. In our work, we call these memories new memories since they are newly formed on each testing day.

### Experimental timeline paradigm

In brief, after confirmation of the stability of the number of neurons with adequate viral expression for calcium imaging and image quality, post-training baseline behaviors in the NOL and CDSD tasks in parallel with neuronal activity recordings (Fig. 1E) in hippocampal panel CA1 were obtained by day -1 (NOL) (Movie 3, the left panel is the calcium activity, right panel the simultaneously recorded behavior) and day 0 (CDSD).

After CDSD task performance on day 0, the 14-day administration of patient or control CSF started (Fig. 1E). Behavior and neuronal activity were recorded on days: [-1, 7, 12, 17, 22, 27, 32, 40, 84] for the NOL task, and on the following days for the CDSD task. A summary of the timeline of the experimental procedures is shown in Fig 1E. We divided the experiment into four phases. The first phase is called “learning” and corresponds to the learning/habituation period where the animals get prepared for the CDSD and NOL tasks. The second phase is called “induction” and lasts from day 0 to 14 where antibodies are infused into the two groups of animals. From day 15 until day 28 we call the early recovery phase and coincide with the observation window of previous studies observation window.^19, 24^ From day 29 until day 85 is the last phase that we call late recovery.

### Statistics

Data from behavioral experiments and calcium imaging recordings were analyzed using two-way ANOVA where “experimental day” and “treatment” were the two independent variables. When ANOVA was significant and one group’s measurements were consistently higher or lower than the other across days we utilized a one-tailed Student’s t-test to determine the difference within individual days. Otherwise, we used a two-tailed Student’s t-test to observe differences within days. We used the memory index (MI, see Supplementary Material document) to quantify the similarity of the same place-cell when the animal is exposed in two different moments to the same environment. To determine the significance of the MI we created bootstrapped surrogate data to determine the [5 to 95] percentile confidence interval. A detailed description of the generation of the surrogate data is explained in the Supplementary Material document.

### Data availability

All data utilized in the figures of this manuscript is provided in the in the public repository: XXXXX. Analysis code in Matlab is uploaded to the webpage (it will be here after acceptance of the manuscript). The computational model and its variations are uploaded in the git-hub repository: https://github.com/BioDatanamics-Lab/NMDAR_Immunoablation_p22_03.

## Results

To determine the underlying neuronal sources of memory deficits during the infusion of NMDAR antibodies, we obtained high-quality calcium imaging on the order of one thousand neurons a day from each of the mice groups: the anti-NMDAR group (n = nine) and control group (n = fourteen). Behaviorally, all animals maintained the motivation to solve the task throughout the 3 months experiment (Suppl. Fig. 4), an important condition to compare memory-based behavioral performance across days.

### NMDAR antibodies induce retrograde and anterograde amnesia

We first established the effect of the antibodies on the animals’ behavior. We compared the two groups while they performed both the CDSD and the NOL tasks, to describe memory deficits in both old and newly formed memories, respectively. We confirmed that at the baseline level, there was no difference in the performance of the CDSD and NOL memory tasks between the anti-NMDAR and control groups (Fig. 2).

**Figure 2:**
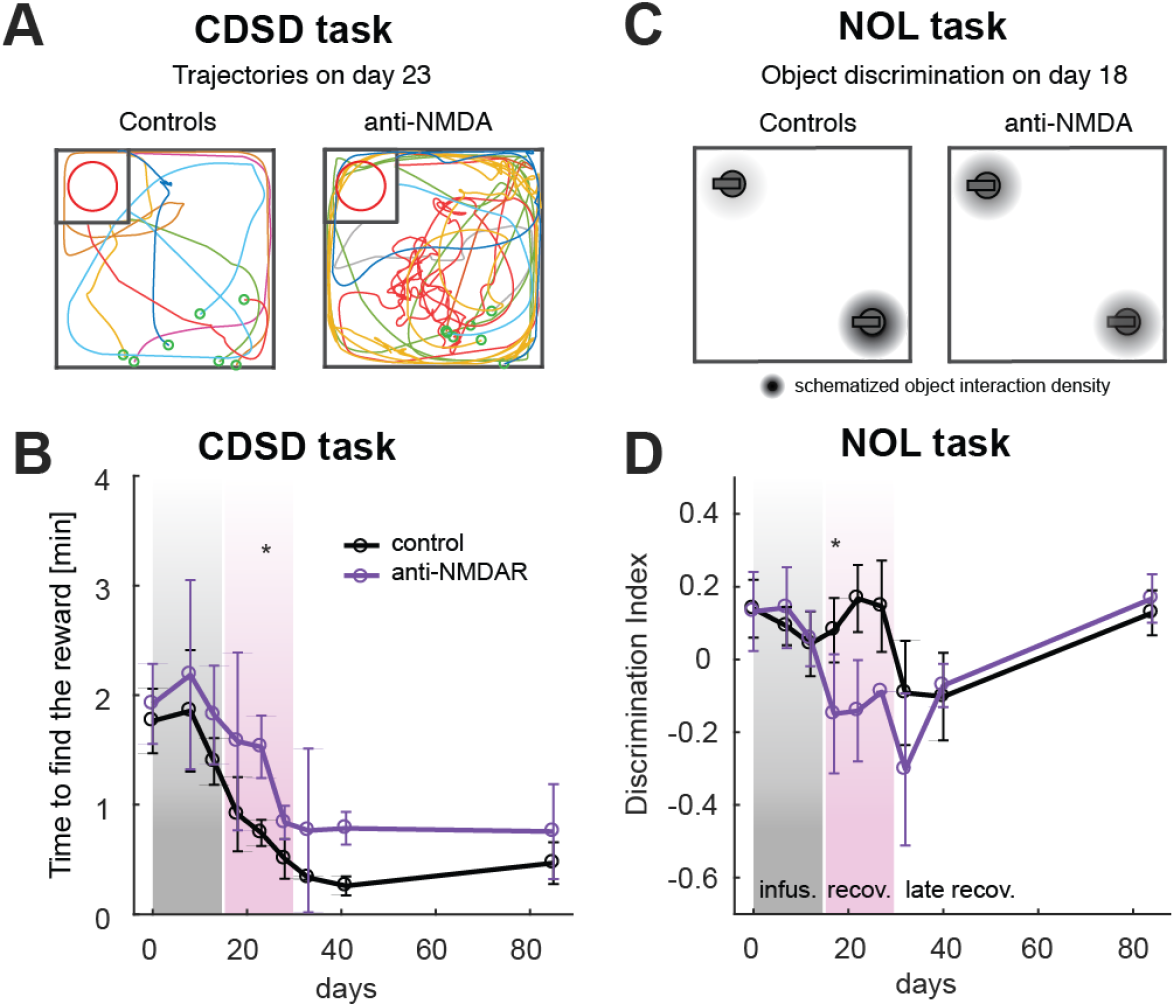
Behavioral memory deficits during and after the infusion of NMDAR antibodies. **(A)** All trajectories from the starting point to the moment the animal engaged with the reward for control animals (left panel) and anti-NMDAR animals (right panel) for experimental day 23 with significant differences between groups’ behavior. **(B)** Behavioral performance of mice in the CDSD task. A shorter time to find the reward indicates better memory of that environment. Anti-NMDAR animals have significantly lower memory on day 23; one-sided 2-sample t-test, p-value=0.048. Note: The time to reach the rewarded location in each environment decreases over time, mainly over the learning phase (from days -4 to 0) and in a smaller amount over the rest of the experiment. (**C**) **Left:** schematic of the behavior of the animal during NOL task performance. Control animals display a bias in the interaction with the object relocated in the test session. **Right:** anti-NDAR animal display no bias (discrimination index close to zero) in the interaction with the object relocated in the test session. (**D**) Behavioral performance of mice in the NOL task. A high discrimination index indicates better formation and maintenance of spatial memory each day. Anti-NMDAR animals have significantly lower 4-hour memory, on day 17; one-sided 2-sample t-test, p-value=0.024.

For the CDSD task, the anti-NMDAR group performed worse than the control group throughout the entire experiment, but the difference was statistically significant (p_value_<0.01) on day 23: 9 days after the end of the antibodies infusion (Fig. 2A-B).

Similarly, in the NOL task, the anti-NMDAR group performed worse than the control group on days 17, 22, 27, and 32 but the difference was significant (p_value_<0.01) only on day 18 as it was previously reported on the same animal model (Fig. 2C-D).^19, 24^ Overall, we found that anti-NMDAR diminished the ability to recall old and new memories behaviorally, where a significant recall impairment occurred during the early stages of the recovery phase.

### Abnormal high neuronal activity during and after the infusion of NMDAR antibodies

To investigate the neuronal basis of the observed behavioral memory dysfunctions, we recorded populations of pyramidal cell neurons in the hippocampal area CA1 (see Methods), while the animals performed the CDSD and the NOL tasks. It has been extensively reported on the existence of spatially tuned excitatory neurons in rodents’ hippocampus called place cells.^25^ These neurons display a preferred spatial location in a closed environment where they tend to fire action potentials more reliably (Fig. 3A). We quantified the place cells’ firing rate as the frequency of calcium events detected in a time window of one second (see Methods) for the neuronal response of the animals moving over the enclosed environment (Fig. 3B). Neurons maintain the same preferred spatial location over time when they are active in the same environment.^26^

**Figure 3:**
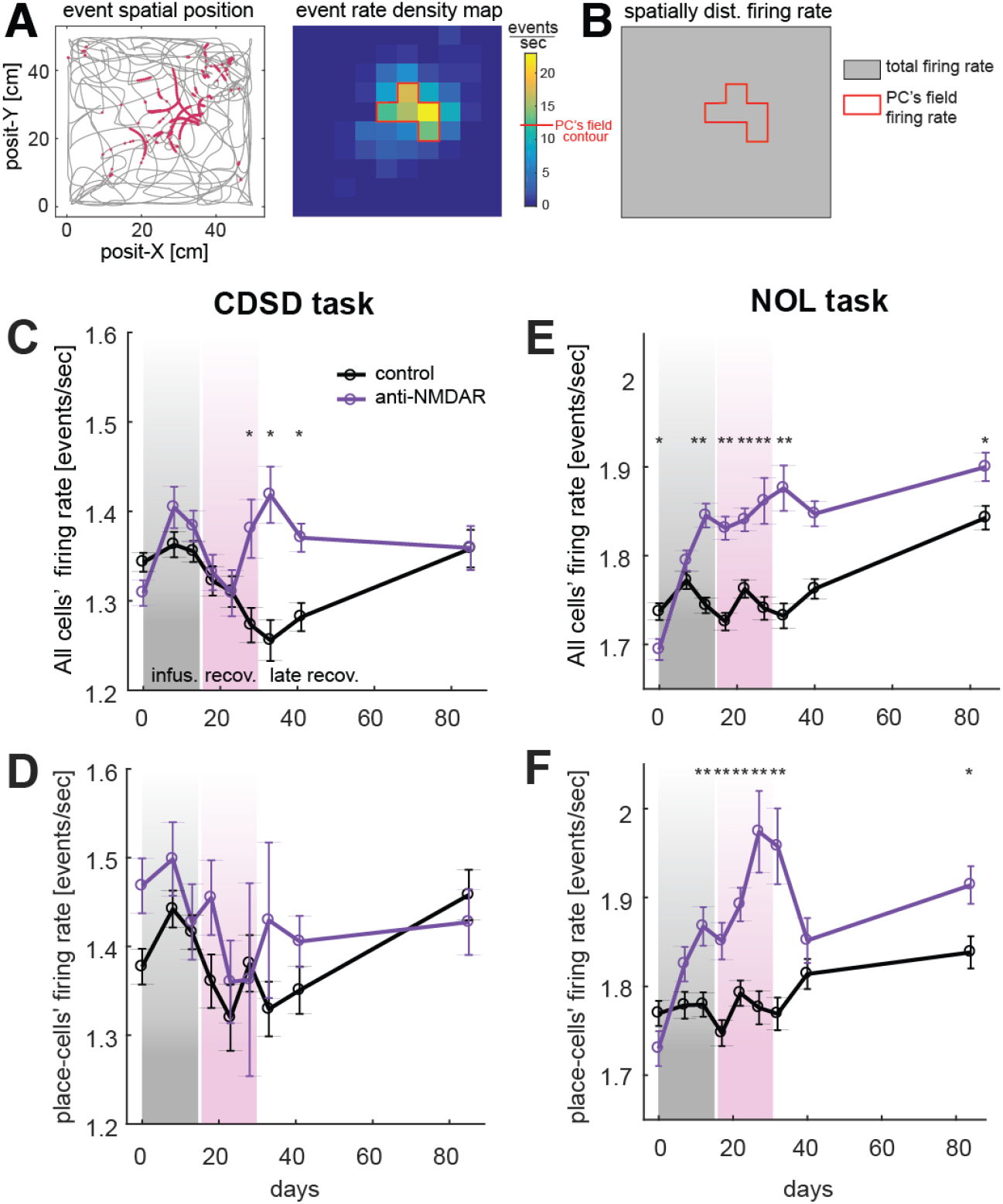
Neuronal hyperactivity during and after the infusion of NMDAR antibodies. **(A)** Example of CA1 place cell. Detected events are plotted at the specific time of the animal’s position. Each event is a burgundy point on the mouse trajectory plot (left). The firing rate for each spatial bin is computed as the number of spikes per second spent on a given spatial bin. The firing rate for different spatial locations is represented with a heat-map (right). (**B**) Description of place cell’s firing rate. Place cell’s field corresponds to all the spatial bins that have equal or more than half the maximum firing rate over the entire environment. The value of half max amplitude is described in the scale for the heat-map in panel A. Place cell firing rate is the mean firing rate over all bins but the largest contribution arrives from the field within the field. (**C**) Mean firing rate of the entire population of cells recorded during the CDSD task across days. The hyperactivating effect of antibodies on neural activity manifests on day 23, and recovers on day 85; two-way ANOVA, p-value(time)=8×10^−4^, p-value(treatment)<10^−4^, p-value(time*treatment)<10^−4^. A one-sided t-test was applied for daily comparison: * = p-value<0.01; **= p-value<0.001. (**D**) Mean firing rate of the place cell subpopulation recorded during the CDSD task. The level of neural activity among place cells in this task was not different between groups; two-way ANOVA, p-value(time)=0.051, p-value(treatment)=0.04, p-value(time*treatment)=0.69 (n.s.). (**E**) Mean firing rate (fr) of the entire population of cells recorded during the NOL task. In the beginning, the cells in the anti-NMDAR group had a lower mean fr than the controls. One-sided t-test, p-value=0.0024. After treatments, fr in anti-NMDAR cells significantly increased and overtook control fr. Two way ANOVA, p-value(time)<10^−4^, p-value(treatment)<10^−4^, p-value(time*treatment)<10^−4^. (**F**) Mean fr of the place cell subpopulation recorded during the NOL task. fr of anti-NMDAR place cells significantly increased due to the presence of antibodies. Two way ANOVA, p-value(time)<10^−4^, p-value(treatment)<10^−4^, p-value(time*treatment)<10^−4^

When we analyzed all the neurons recorded in CA1, we found that in the CDSD task, during the antibodies infusion period, there was no significant difference in firing rate activity between experimental groups (Fig. 3C). Instead, in the early recovery period, we measured a decrease in the firing rate in both the anti-NMDAR and control groups. Notice that a similar decrease in firing rate has been previously reported in area CA1 as the product of the habituation to the behavioral boxes.^27^ After an initial decrease in firing rate in both groups, there was a significant increase in the firing rate in the anti-NMDAR relative to the control group, during the early recovery period. This difference in firing rate between groups disappeared ∼70 days after the antibodies infusion ended (day 85). When the firing rate of only place cells was considered, the difference between groups was not significant (Fig. 3D). For the NOL test, the anti-NMDAR group firing rate was larger than for controls animals, and increased significantly around day 10 of the antibodies infusion period, and remained higher even ∼70 days after antibodies infusion ended (Fig. 3E). The firing rate of only the place cells also displayed a significant increase compared to the control groups (Fig. 3F) in the NOL task.

Together, these results indicate that the effect of antibodies is to generate neuronal hyperactivity in the entire population of CA1 neurons, but the effect is stronger in newly learned memories than in old memories formed previous to the antibodies infusion.

### Degradation of place cells’ spatially modulated responses due to NMDAR antibodies

It has been well established that neurons that respond to a particular set of stimuli (i.e. preferred stimuli) allow the brain to decode the information encoded in the neuronal population response to that stimulus (for review see ^28^). However, if the amplitude of the response modulation induced by the stimulation is smaller than the response variability over different stimulations, the information encoded by the neuron can drastically be diminished or even abolished (Suppl. Fig. 5). A decrease in total signal to noise ratio (SNR) is proportional to a decrease in total information encoded by the neurons.^29^ In principle, the increase in total firing rate observed in our data (Fig. 3C-E) could increase the spatially modulated responses (i.e. signal increase), but could also increase the spatial-independent neuronal response (i.e. noise increase), potentially leading to a net decrease in the SNR depending on the sizes of each of the components.

We hypothesized that the significant increase in the firing rate of the anti-NMDAR group relative to controls could cause a reduction of spatial information leading to the memory deficits observed behaviorally. To test this hypothesis we first investigated changes in the activity of place cells during and after the infusion of antibodies. Reduction in spatial information in place cells leads to a poorer brain representation of space that could affect navigation and memory.^30^ To be able to measure experimentally changes in neurons’ preference for spatial locations with high statistical power, we quantified the effect of an increase in total firing rate measuring the change in the ratio between the place cell’s mean firing rate in the inside versus the outside field for each given cell on each recording session (Fig. 4A, Suppl. Fig. 5). To this end, the place cell’s field is defined here as the set of bins where the firing rate is larger than half maximum rate over the entire environment (Fig. 4A & Supplementary Material). We then call the firing rate inside-field the average firing rate for all the spatial bins within the field, and the firing rate outside-field is the average firing rate over the rest of the bins that tessellate the environment. If the ratio of firing rate inside/outside decreases, then information diminishes (for a constant amount of noise). In this section, we solely quantified the change in the inside/outside place cell response ratio to gain insight about the effect of the global increase in firing rate due to the infusion of antibodies. Understanding the effect of noise will be the focus of the following section.

**Figure 4:**
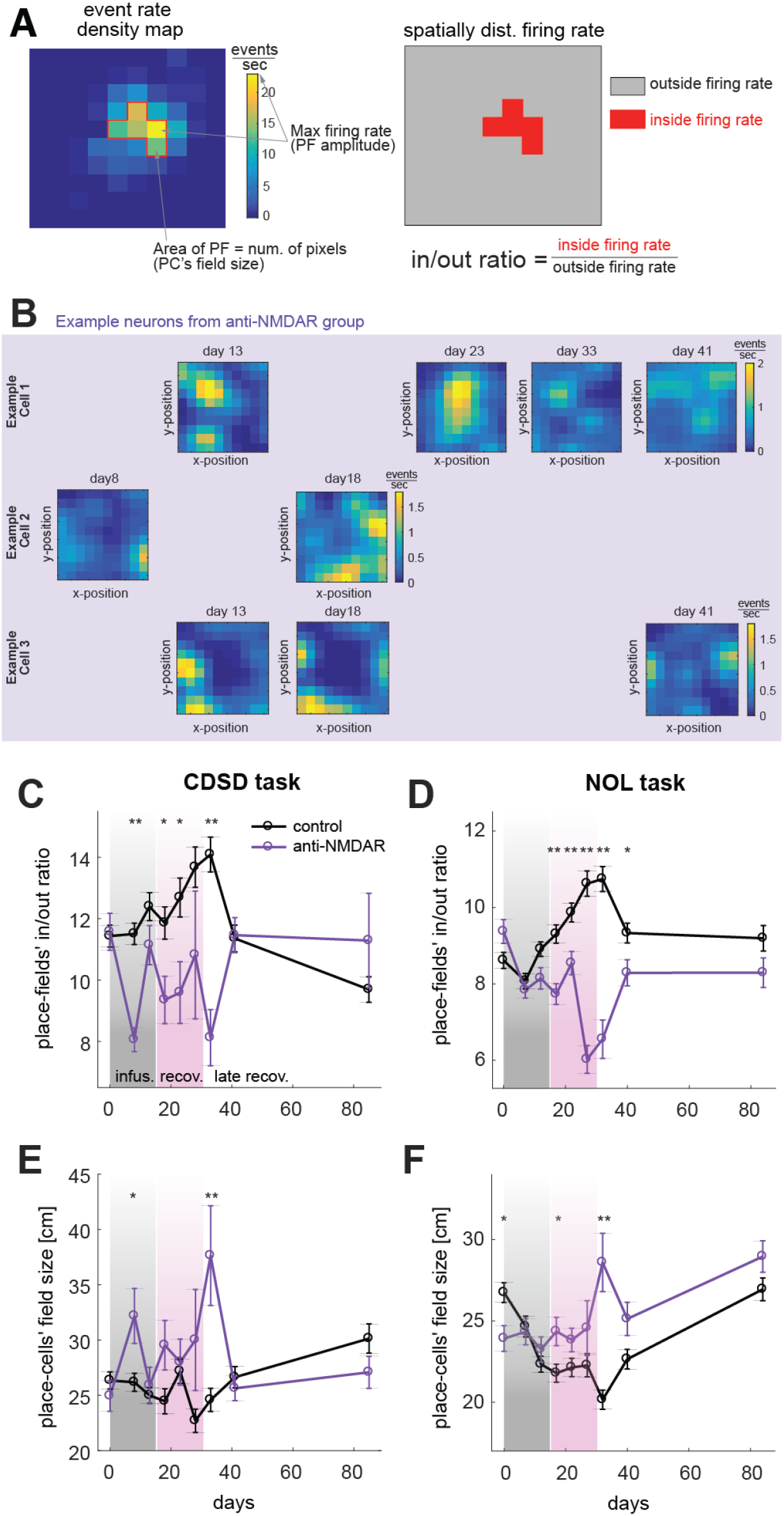
Decrease in neuronal spatial tuning due to the infusion of NMDAR antibodies. (**A**) Description of place cells parameters: amplitude, in/out ratio, and field size. (**B**) Three examples of neurons’ place cell’s fields over different sessions of the experiment for the anti-NMDAR group. We can measure place cells over time because with our technique we can follow the same neurons for ∼3 months. Missing spots correspond to neurons inactivating for certain sessions, a phenomenon called turn-over (Ziv et al 2013). place cell’s fields change over time in the anti-NMDAR group, for example, increasing activity in the outside-field and the inside-field becoming more distributed over the environment. (**C**) The ratio of control place-fields activity (inside/outside) in the CDSD task was higher than anti-NMDARs due to antibodies effects. Two way ANOVA, p-value(time)=0.18 (n.s.), p-value(treatment)<10^−4^, p-value(time*treatment)=4×10^−4^. A one-sided t-test was applied for daily comparison: * = p-value<0.01; **= p-value<0.001. (**D**) The ratio of control place-fields activity (inside/outside) in the NOL task was higher than anti-NMDARs. Two way ANOVA, p-value(time)=7×10^−4^, p-value(treatment)<10^−4^, p-value(time*treatment)<10^−4^. **(E)** The average place-field size of place cells in the CDSD task. Place-field of anti-NMDAR place cell was significantly enlarged (purple) compared to controls (black) due to the effect of antibodies. Two way ANOVA, p-value(time)=0.014, p-value(treatment)=2×10^−4^, p-value(time*treatment)=1.4×10^−4^. (**F**) The average place-field size of place cells in the NOL task. Control place cells (black) had on average larger place cells fields in the beginning; one-sided t-test, p-value=0.0026, but their place fields gradually shrank throughout the experiment days and remained significantly smaller than anti-NMDAR place fields. Two way ANOVA, p-value(time)<10^−4^, p-value(treatment)<10^−4^, p-value(time*treatment)<10^−4^.

We identified that the decrease in the inside/outside ratio is primarily due to spreading cell firing outside the main place cell’s field (Fig. 4B), and therefore neuronal spatial tuning decreases during the effect of NMDAR antibodies. We measured that the inside/outside ratio was significantly lower for the anti-NMDA group from days 8 to 41 in the CDSD test and from days 17 to 40 in the NOL tests (Fig. 4C-D).

These results demonstrate that the increase in firing rate observed in the hippocampal neuronal population during and after infusion of NMDAR antibodies is associated with a reduction of the place cell’s spatially modulated responses for both new and old memories. Specifically, we show that changes in the inside/outside ratio coexist with an increase in place cells’ field size in the anti-NMDAR group relative to controls. Even though the increase in field size was only significant on days 8 and 33 of the experiment in the CDSD task, and on days 17 and 32 for the NOL task, the trend throughout the experiment after the start of the antibodies infusion (Fig. 4E-F) was the same. This observation is consistent with a decrease in spatial information due to the presence of antibodies. In the next section, we extend our analysis by taking into account the effect of the noise.

### CA1 spatial information is reduced due to NMDAR antibodies

Our results demonstrated that the reduction in NMDAR leads to an increase in neuronal activity and changes in place cell’s field shapes. As mentioned above, estimating the total effect of such place cell’s field changes requires assessing the contribution of the noise. Here we will determine spatial information including the effect of noise to establish how antibodies can affect cognitive processes like navigation and spatial memory. To ultimately quantify how antibodies changed the spatial information encoded in the CA1 neuronal activity, we computed the signal-to-noise ratio (SNR, see Supplementary Methods) of the entire population of neurons and not only of the place cells as in the previous section. Spatial information is carried by place cells but can also be carried by mixed-selective neurons.^31,32^ For this reason, we computed SNR for an optimized spatial and temporal discretization of the calcium events (see Suppl. Methods). Because we wanted to observe the percentage changes in spatial information at different time points relative to the state prior to the presence of the antibodies, we normalized the SNR for the anti-NMDAR and control groups relative to its baseline values for each group independently.

For the CDSD task, we found that the SNR of the control group continuously increased during the experiment and decayed at the end (days 41 and 85 in Fig. 5A). In contrast, in the anti-NMDAR group the SNR decreased during and after the infusion of antibodies, and on days 41 and 85, it had recovered to levels closer to baseline.

**Figure 5:**
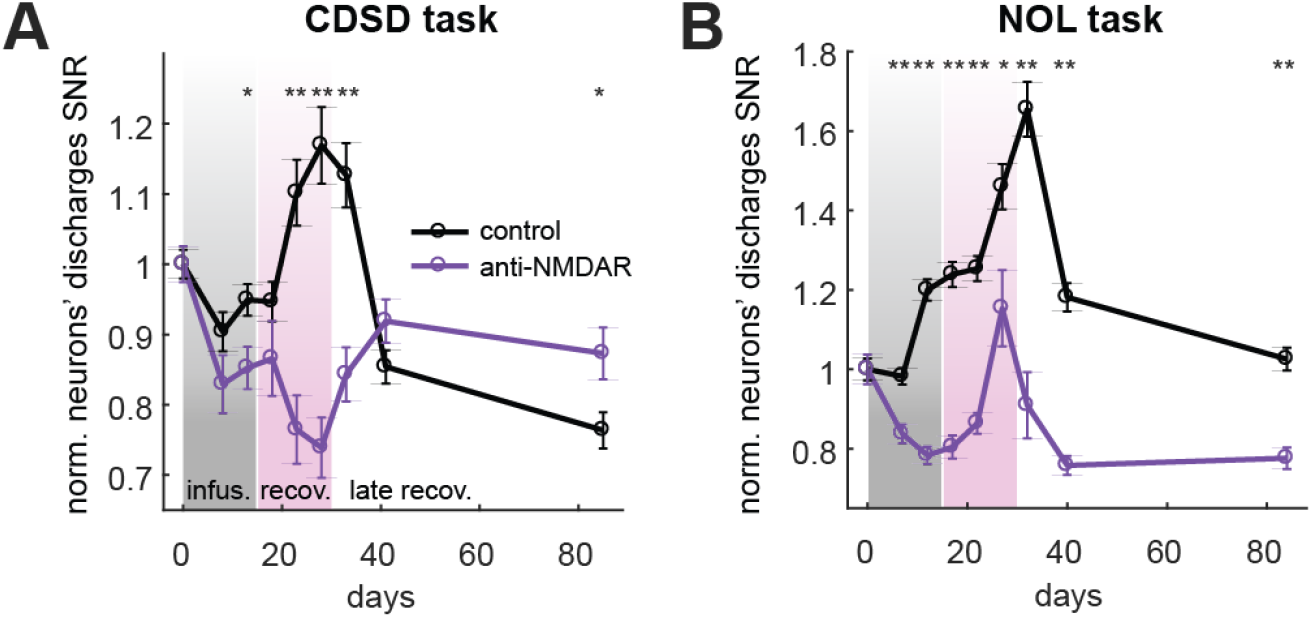
Degradation of spatial information due to NMDAR antibodies. (**A**) Spatial information of control cells was significantly higher than anti-NMDAr cells in the CDSD task. Two way ANOVA, p-value(time)<10^−4^, p-value(treatment)<10^−4^, p-value(time*treatment)<10^−4^. A one-sided t-test was applied for daily comparison: * = p-value<0.01; **= p-value<0.001. **(B**) Spatial information of control cells was significantly higher than anti-NMDAR cells in the NOL task. Two way ANOVA, p-value(time)<10^−4^, p-value(treatment)<10^−4^, p-value(time*treatment)<10^−4^.

In the NOL task, we found that the SNR of the control group increased from baseline to day 32, and decreased during the late recovery phase, when the time intervals between tests are larger (Fig. 5B). On the other hand, the SNR of the anti-NMDAR group decreased during the induction phase and then increased during the early recovery. During the late recovery phase, the SNR did not recover to its baseline values even 70 days after the infusion of antibodies had stopped.

Together, these results show that in our mouse model, the hippocampal spatial information is lower in anti-NMDAR groups during and after the infusion of antibodies for old and newly formed memories.

### Lack of place cell’s temporal stability can explain NMDAR antibodies evoked retrograde and anterograde amnesia

We have shown so far that animals in the anti-NMDAR group display higher firing rates which led to a decrease in CA1 neurons’ spatial information. The hippocampal research field utilizes the stability of the place cell’s fields over time as an indicator of spatial memory.^11^ Thus, it is reasonable to hypothesize that if spatial information decreases, then the temporal stability of the place cells also decreases, reducing the total memory encoded in the neuronal population. To measure the temporal stability of the place cell’s field, we created the memory index (MI), a measurement of similarity between a place cell’s field over time (see Suppl. Methods).

Over the time scale of minutes, we did not find consistent MI differences between the anti-NMDAR and control groups in either of the two tasks (see Suppl. Fig. 6), demonstrating that during exposure to NMDAR antibodies place cells’ fields are stable over short time scales. To examine the effects over a time scale of hours we analyzed data from the NOL task since the difference between the training and the testing sessions was ∼4 hours. This analysis showed that the control group had a higher MI than the anti-NMDAR group for almost every tested day of the experiment. In particular, the difference was significant on days 7, 22, and 32 (Fig. 6A), and the difference disappeared by day 84.

**Figure 6:**
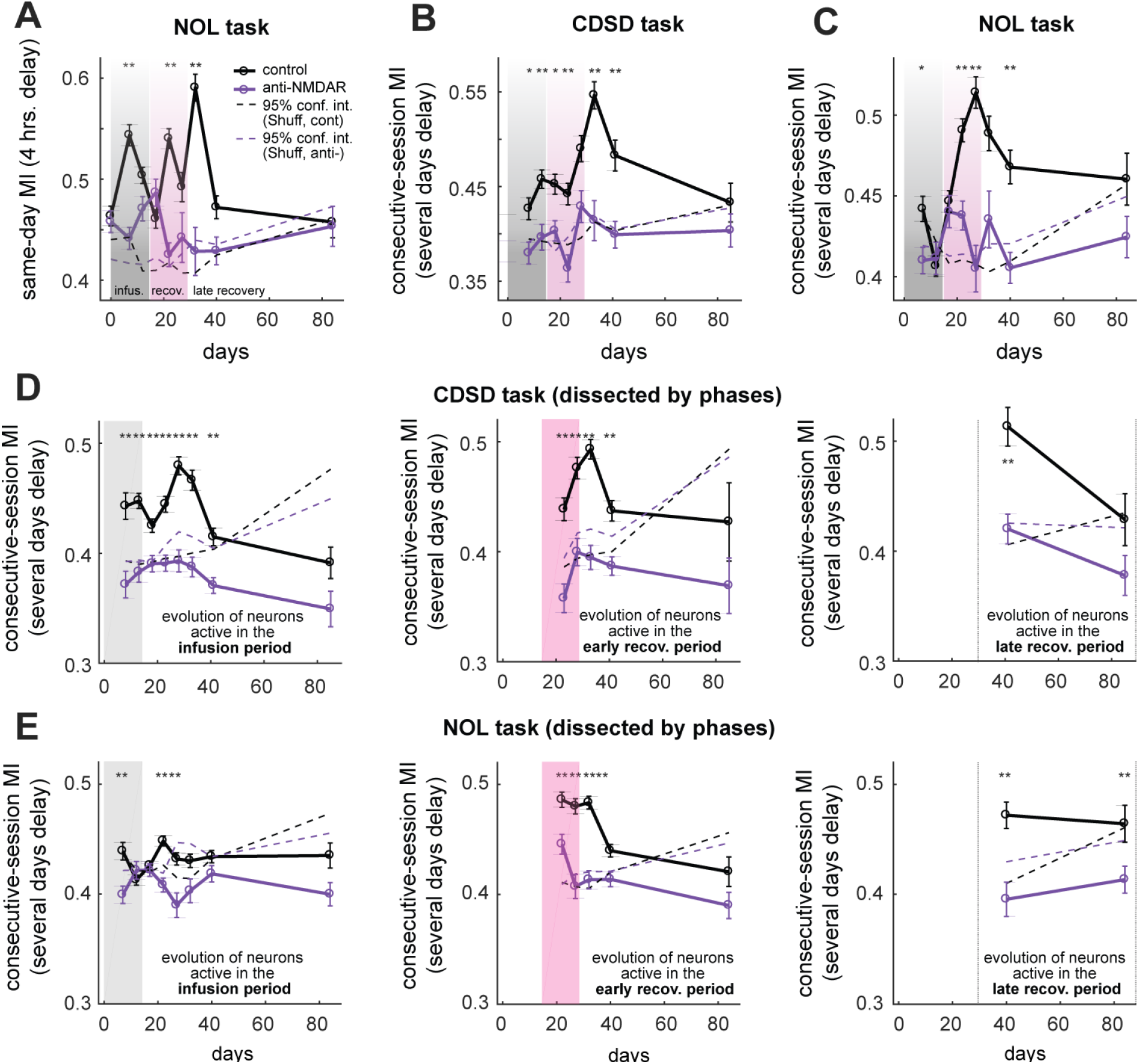
Degradation of old and new memories due to the presence of NMDAR Abs. In all panels, the dashed lines show the significance threshold of MI with 95% confidence interval. When solid curves are above dashed lines, place cells’ memory is significantly higher than chance. **(A)** Mean MI for cells that were active in both training and recall sessions of given NOL days. Mean MI became higher among control cells than anti-NMDAR cells and recovered on day 84. Two way ANOVA, p-value(time)=8×10^−4^, p-value(treatment)<10^−4^, p-value(time*treatment)<10^−4^. A one-sided t-test was applied for daily comparison: * = p-value<0.01; ** = p-value<0.001. **(B)** Mean MI for cells that were active in two consecutive days of the CDSD task, within an identical environment (i.e. Env.A-day(n) and Env.A-day(n+1), or Env.B-day(n) and Env.B-day(n+1)). Mean MI became higher among control cells than anti-NMDAR cells and recovered on day 85. Two way ANOVA, p-value(time)<10^−4^, p-value(treatment)<10^−4^, p-value(time*treatment)=0.13 (n.s.). **(C)** Mean MI for cells that were active on two consecutive days of the NOL task, within identical sessions (i.e. training-day(n) and training-day(n+1), or testing-day(n) and testing-day(n+1)). Mean MI became higher among control cells than anti-NMDAR cells and recovered on day 84. Two way ANOVA, p-value(time)<10^−4^, p-value(treatment)<10^−4^, p-value(time*treatment)<10^−4^. **(D)** Left panel: longitudinal tracking of the MI for the cells which became active during the induction phase. Two way ANOVA, p-value(time)<10^−4^, p-value(treatment)<10^−4^, p-value(time*treatment)=0.07 (n.s.). Middle panel: longitudinal tracking of the MI for the cells which became active during the early recovery phase. Two way ANOVA, p-value(time)=7×10^−4^, p-value(treatment)<10^−4^, p-value(time*treatment)=0.22 (n.s.). Right panel: longitudinal tracking of the MI for the cells which became active during the late recovery phase. Two way ANOVA, p-value(time)=0.003, p-value(treatment)=8×10^−4^, p-value(time*treatment)=0.32:(n.s.). In all three panels, control cells’ MI is always significantly higher than anti-NMDARs. **(E)** Left panel: Tracking MI for the cells which became active during the induction phase and one or more of the future days. Two way ANOVA, p-value(time)=0.3 (n.s.), p-value(treatment)<10^−4^, p-value(time*treatment)=5×10^−4^. Middle panel: Tracking MI for the cells which became active during the early recovery phase and one or more of the future days. Two way ANOVA, p-value(time)<10^−4^, p-value(treatment)<10^−4^, p-value(time*treatment)=0.01. Right Panel: Tracking MI for the cells which became active during the late recovery phase and one or more of the future days. Two way ANOVA, p-value(time)=0.76 (n.s.), p-value(treatment)=2×10^−4^, p-value(time*treatment)=0.45 (n.s.).

MI is a measurement that depends on the size of the place cell’s field in relation to the size of the behavioral environment. To give a reference level for the amplitude of the MI we compared it with the MI obtained by measuring the similarity between two random place cell’s field from two consecutive sessions, and then we repeated this procedure over all possible pairs of neurons (see the bootstrapping procedure in Methods). This procedure provides a [5-95] percentile confidence interval for the significant MIs. Thus, MI under the 95 percentile of the surrogate data’s confidence interval displayed a non-significant 4hrs place cell stability (Fig. 6 dashed lines; see also Supplementary Methods). The result in figure 6A is consistent with a decrease of NMDAR affecting the formation of new memories on the time scale of hours, as previously reported using NMDAR antagonists.^10,11^ Therefore, during the late recovery phase, it will be difficult to record the same pattern of neuronal activity over time, which could directly or indirectly affect hippocampal-encoded memory.

We then extended our analysis and studied how the antibodies affected the maintenance of long-term memories for several days, a time scale that is not addressed in human patients’ neuropsychological assessments. We looked at differences for days intervals of 5, 10, and 45 days. In this analysis the MI plotted on the figure corresponds to the value from the latest point of the time interval: for example, MI on day 8 displays the similarity of the place cells that were active on days 0 and 8. We observed a significantly lower memory strength in the anti-NMDAR group relative to the control group for the time intervals ending on day 8 to 41 in the CDSD task (Fig. 6B); and 7, 22, 27, and 40 for the NOL task (Fig. 6C). Importantly, the control group displayed strong long-term memories for most consecutive-sessions intervals throughout the experiment. In contrast, the MI in the anti-NMDAR group remained close to the chance level, showing negligible values of hippocampal neuronal correlates of long-term memory throughout the experiment.

Overall, these findings show that the neuronal correlate of new memories degrades over periods of 4 hrs due to the presence of NMDAR antibodies. After infusion of antibodies, the memory encoded in the CA1 neurons’ activity was negligible or inexistent for the majority of the time intervals for old and new memories.

### The long-lasting effect of NMDAR antibodies on long-term place cells stability

We have demonstrated that the preferred spatial neuronal response location (i.e. place cells’ field) is unstable across days when the NMDAR antibodies are present, supporting the concept of a neuronal malfunction explanation for retrograde and anterograde amnesia observed in our behavioral experiments. In our data so far, we have looked at the phenomenon of memory in a rather static framework from the point of view of individual time intervals between minutes, hours, and days. However, memories are dynamic mnemonic elements that change every time we are exposed to similar situations and contexts.^33^ Therefore, we then used our data set to examine how the neuronal correlates of memories (i.e. place cells’ stability) evolved and investigated how the presence of NMDAR antibodies affects their evolution.

To quantify the effect of NMDAR antibodies on the temporal dynamics of memory formation, we studied how memories generated at different time points in our experiment evolved during and after the infusion to the antibodies. Most of the theories on memory formation^34^ agree on interpreting memory strength as a dynamic variable that increases every time a subject is exposed to the same stimulus in the same context.^14^ Without a novel exposition, memory strength decays over time to a lower value, vanishing in some cases (i.e. passive forgetting). If the frequency of exposition to the stimuli is high, then one will expect a net increase in memory (i.e. massed/spaced training). Instead, if the frequency is sufficiently low, then each exposition could lead to a constant amplitude memory strength, which is almost unaffected by the value of previous expositions (i.e. maintenance or spaced re-learning). For lower frequencies, the memory strength could decay to zero.

In the CDSC task, the control group showed first a period of increasing MI for each of the phases, and then a natural decay, significantly decreasing the place cells’ stability (Fig. 6D). The later the phase, the higher the maximum MI reached, as expected by the effect of training on previous sessions. In contrast, the anti-NMDAR group had no significant MI over the entire experiment (solid line below dashed line), even for place cells that were activated for the first time two weeks after the infusion of the antibodies (Fig. 6D, right panel).

In the NOL task, the only significant MI that was detected for the anti-NMDAR group occurred at the beginning of the recovery phase, as a result of learning during the previous phase when the levels of antibodies are still low (Fig. 6E). For the rest of the experimental days, the MI was below chance describing the strong effect of antibodies on newly formed memories. This last result provides a potential explanation for why the recovery of NMDARe patients is slow and can last from months to years.^6,7^

Together, these results demonstrate that the evolution of MI for new memories leads to severe instability of place cells’ fields due to the infusion of antibodies. The MI during the recall of memories learned prior to the antibodies infusion is prominently reduced, displaying a neuronal abnormal behavior consistent with retrograde amnesia.

### Main factors that could explain neuronal activity alterations induced by NMDAR antibodies

Changes in NMDAR densities have been mainly associated with changes in synaptic plasticity such as long term-potentiation^35^ between a presynaptic and a postsynaptic cell, which can indirectly affect spatial information by shaping place cells responses.^36,37^ Less known is the role of the NMDAR-mediated currents in balancing the excitatory and inhibitory inputs that dictate the neurons’ firing rate.^38,39^ To address this, we built a simplified hippocampal network model that allows the study of how a systemic reduction of an excitatory current such as the NMDAR-mediated current, leads to a global increase in neuronal activity at the network level.

Our model (Fig. 7A, Suppl. Methods: Annex I) was designed to comply with a minimum, but a necessary number of premises. It consists of a feed-forward network mimicking the Schaffer collateral pathway (see detailed description below), an integral part of hippocampal memory circuitry. The model includes excitatory and inhibitory neurons that change their firing rate depending on the position of the animal in the space, which in this simple model is a linear track. Neurons include an excitatory current (interpreted as combining AMPA and NMDA components) that has the same amplitude parameter in both excitatory and inhibitory cells.^40^ In our simplified model, each NMDA current term grows linearly with the neuron’s firing rate, for excitatory and for inhibitory neurons. Simpler models with constant amplitude of NMDA did not provide the neuronal behavior observed in our data (Suppl. Fig. 7). Models with the dependence of the NMDA currents on the synaptic inputs would be more realistic, but would provide lower tractability of the relevant variables.

**Figure 7:**
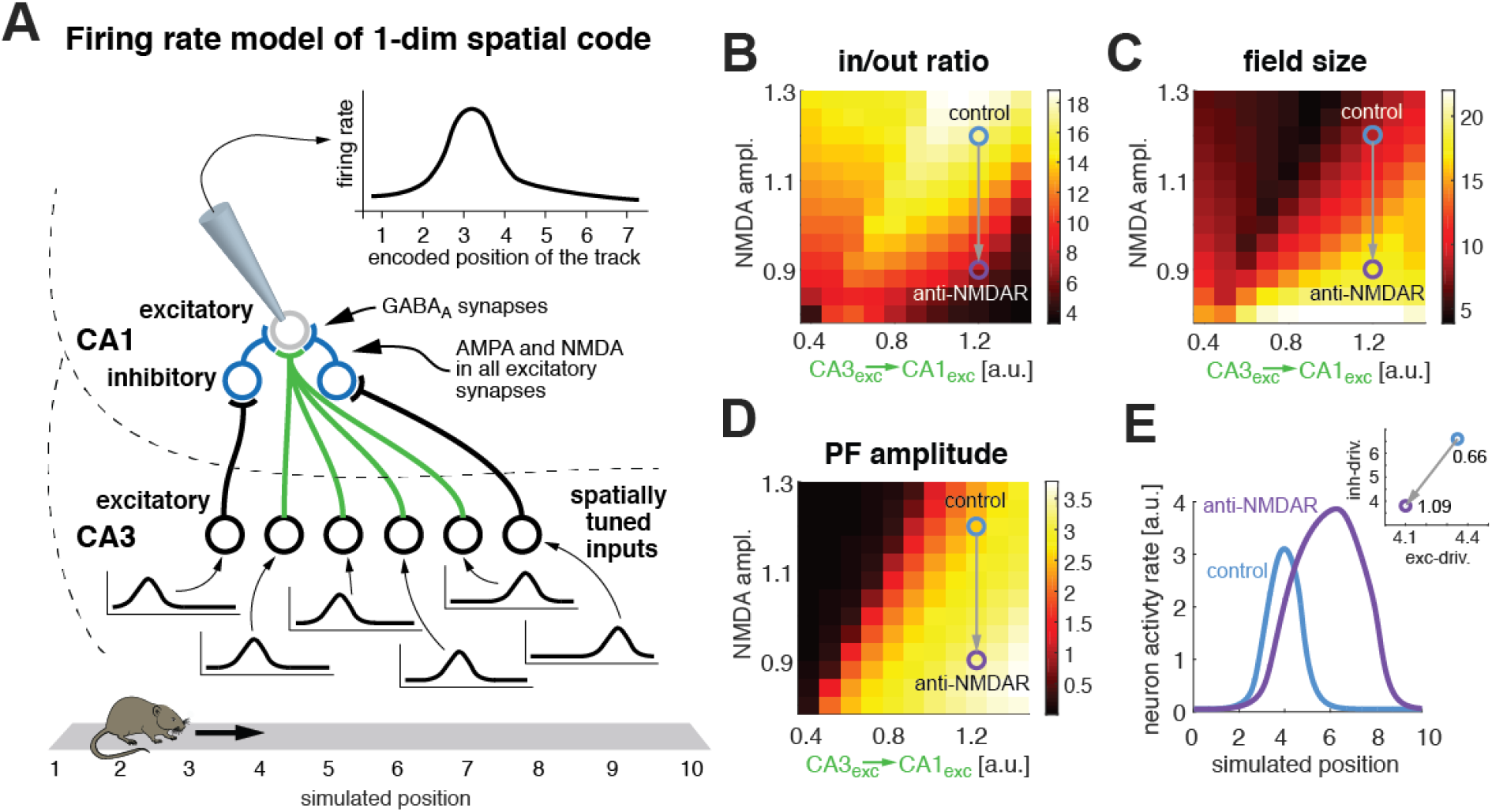
Computational model of area CA1 affected by the reduction of NMDAR. **(A**) Schematic of the model’s connectivity between layers. The model is a feed-forward network of rate-based neurons that include AMPA and NMDA activity-dependent currents. CA3 neurons are spatially modulated, resembling place cells that provide tuning to a CA1 via a gaussian shape connectivity structure. The model mimics the response of a single cell when the animal walks in a straight line. (**B**) Heat map for the Input/Output ratio. Parameter search of NMDA current amplitude factor and the connectivity factor between excitatory cells in CA3 and the excitatory cell in CA1. The optimal parameter set was found over a high dimensional space of connectivity parameters, where we obtained the two main factors by manual inspection. **(C**) Place cell’s field size heatmap for the same parameter search space as in B. (**D**) Place cell’s field amplitude heatmap for the same parameter search space as in B. (**E**) Place cell responses for the control network (blue trace) and the NMDAR antibodies network (purple trace). Inset: excitatory vs. inhibitory driving force at the maximum amplitude point during the place cell response in the CA1 neuron. The balance of excitation inhibition changes from a ratio exc/inh of 0.61 in the control to 0.98 in the anti-NMDA group showing that the effect of NMDAR is stronger on the inhibitory inputs.

Specifically, our model includes three layers: i) CA3 excitatory cells (CA3-exc), that target ii) CA1 inhibitory cells (CA1-inh), and iii) a single readout excitatory neuron in CA1-exc that exemplifies our average recorded neuron (see, Supplementary Methods: Annex I). Each CA3-exc generated a place-tuned response that fed the CA1 layer (Fig. 7A). We measured the output of the CA1-exc as a readout of the effect of the presence of NMDAR antibodies in the network. By construction, we have balanced the network so that the individual inhibitory input to the CA1-exc neurons is ∼5 times larger than the excitatory inputs from CA3-exc. This condition was necessary to generate a single place field, by adding the input from twenty excitatory neurons in CA3 and one inhibitory neuron in CA1. The ratio between excitation and inhibition synaptic strength is similar to the one observed experimentally.^41^

By decreasing the amplitude of NMDAR-mediated currents, mimicking the effect of the antibodies, we reduced the excitatory input to both CA3-exc cells and CA1-inh cells by equal amounts. With this simple manipulation in the model we reproduced the general behavior of reduction of inside/outside place cell’s field ratio (Fig. 7B), the increase in place cell’s field size (Fig. 7C), and the rise in place cell’s field amplitude (Fig. 7D) observed in our experiments. The changes in the place cell’s field shape due to the decrease in NMDAR can be visualized from the model’s output neuron activity CA1-exc (Fig. 7E). Notice that the inside/outside place cell’s field ratio is taken between the spatial bins with larger half-max amplitude and the ones with less of the half-max amplitude, correspondingly.

Our simplified model displays the main aspects of our experimental observations derived from the NMDAR decrease. It also provides a key observation regarding the paradoxical phenomenon present in our data, that is, that a systemic reduction in NMDAR leads to a reduction chiefly in the inhibitory driving force, which leads to, or induces, global hyperactivity observed in our experimental recordings. The reduction in the NMDAR-mediated currents in CA3 and CA1 neurons causes a decrease in the E-I balance ratio (Fig. 7E, Inset).

## Discussion

Previous animal models of immunoablation of NMDAR have substantially contributed to understand the molecular mechanisms of anti-NMDARe and to establish the antibody pathogenicity regarding the cognitive and synaptic alterations of the disease.^9,24^ However, these studies were unable to establish a relationship between antibody effects at the molecular level and the changes at neuronal network level, which requires the recording of neuronal activity while animals perform cognitively demanding tasks. By being able to carry out this type of experiment, we provide a potential explanation for the antibody-mediated memory deficits based on the disruption of neuronal activity in hippocampal CA1 cells.

Our findings show that the antibodies cause hippocampal neuron hyperactivity, which leads to degradation of the spatial information carried by cells in CA1. Increase in neuronal activity does not necessarily lead to a decrease in information, but we observed that the hyperactivity led to an increase of the noise and decrease in the signal components, creating a net decrease in SNR, which translates into a decrease in information. This is consistent with previous studies showing that CA1 neuronal activity degradation associated with lower performance in spatial navigation,^42,43^ and it has also been reported in several animal models of cognitive alterations.^44-47^ In addition, we observed a reduction of neuronal response stability over time – a measurement of memory strength.^48,49^ The current findings provide evidence of retrograde amnesia, a cognitive dysfunction that has not been explored in anti-NMDARe patients but has been observed in other synaptopathies.^50^ In our model we found that under the effect of patients’ antibodies, the neuronal responses stability over time diminishes more for newly formed memories than for memories learned prior to the administration of the antibodies. These results are consistent with the role of NMDAR in memory consolidation, where antibodies would affect less the memories that have been already consolidated than those that are newly formed. Notably, the antibody-mediated changes at the neuronal network level lasted much longer than the synaptic NMDAR density recovery consistently reported in this animal model (i.e, day 24).^9^ Given that in this model there are minimal or absent inflammatory changes,^9^ the antibody-mediated effects such as reduction of NMDARs and the prolonged disruption of network function demonstrated here, seem to be sufficient for the protracted memory and cognitive dysfunction observed in patients with this disease.

It is well known the role of NMDAR in the formation of memories at the single cell level, but why can an acute decrease in its density lead to long-term abnormal behavior for the entire hippocampal neuronal network? When NMDAR is reduced in our computational model we observe an hyperactivity state and a degradation of the CA1 spatial information due to change in the balance of the excitatory and inhibitory inputs to the CA1 output layer. This excitation/inhibition unbalance has been suggested in prior studies using different approaches. For example, using an animal model of passive antibody transfer similar to the model used in our work,^51^ showed that NMDAR antibodies caused an ictogenesis state similar to that occurring in patients. With a computational model, they showed that the transition to an ictogenesis was best explained by changes in synaptic weights that pushed excitation/inhibition out of balance. Ceanga et al.^52^ used in-vitro recordings and a computational model to demonstrate that changes in excitation/inhibition balance caused by lower AMPAR strength and faster GABAA-receptor kinetics, mainly accounted for the abnormal EEG oscillations. Therefore, we provide here a different set of studies focused on memory and behavior, in which we were able to link the excitation/inhibition unbalance with alterations of these tasks.

The reasons why the antibody-mediated alteration of neuronal activity lasted longer (i.e., ≥70 days) than the alterations observed with the behavioral tests (18-23 days for NOL and CDSD) are likely related to different levels of sensitivity of these paradigms: the sensitivity of neuronal recordings is likely greater than the sensitivity of the indicated behavioral paradigms. From where we stand now, it seems possible that the development of more demanding behavioral tasks in mice may reveal long-term behavioral effects that will align better with the neuronal hyperactivity and degradation of memory and spatial information. Another task for the future is to determine how recent and newly learned spatial locations affect the spatial navigation of patients with anti-NMDARe.

Overall, the current study provides a mechanism that likely underlies the long-term effects of antibody mediated reduction or NMDAR. The animal model can now be used for additional investigations on the role of NMDAR in memory maintenance and recall as well as for novel approaches (i.e., allosteric modulators of NMDA or AMPA receptors) to treat the long-term memory and behavioral alterations of the disease.^24,52^

## Abbreviations

(NMDAR): NMDA receptor
(anti-NMDARe): NMDA receptor encephalitis
(CSF): cerebrospinal fluid
(NOL): novel object location
(CDSD): context-dependent spatial discrimination

## Acknowledgments

We are thankful for the insightful comments and discussions from Myrna Rosenfeld, Helena Ariño, and Albert Compte. PEJ and JD designed the experiment and obtained the funding. PP-M and PEJ performed the calcium imaging recordings and the behavioral tests. PEJ performed all surgeries. APZ, PP-M, PEJ have performed mice behavioral analysis. APZ performed the calcium imaging analysis. DTC wrote the calcium imaging preprocessing code. HGR designed and wrote the computational model code. PEJ analyzed the model output. PEJ wrote the manuscript and all the authors contributed with the writing procedure.

## Funding

We all gratefully acknowledge the funding support of this work. Fundació CELLEX supported PP-M, DPT, JD, and PEJ; Marie Curie Fellowship FP7-PEOPLE-2013-IIF (ISOLM; Ref#: 627457) supported DPT and PEJ; APZ, PP-M and PEJ were supported by the grant PID2019-110427RB-I00, from the Ministry of Science, Innovation and Universities. “La Caixa” Foundation (ID 100010434 under agreement LCF/PR/HR17/52150001) supported APZ, PEJ and JD; JD was supported by Instituto Carlos III (Proyectos Integrados de Excelencia 16/00014 and FIS PI20/00197) the Edmond J Safra Foundation; HGR was supported by the USA National Science Foundation grant IOS-2002863.

## Competing interests

The authors declare no conflicts of interest related to this work.

## Supplementary material

Supplementary material is available at Brain online

## Notes

### Competing Interest Statement

The authors have declared no competing interest.

### Summary of Updates

New version with figures within the main text

