## Supplementary material for "Anti-NMDAR encephalitis antibodies cause long-lasting degradation of the hippocampal neural representation of memory": Suplementary material (from another version of this preprint: version 1, 2022-11-25)

**SUPPLEMENTARY INFORMATION**

1. **Mouse model of antibody-mediated NMDAR Ab transfer**
2. **Calcium imaging + intraventricular infusion preparation**
3. **Calcium activity recordings**
4. **Behavioral Analysis**
5. **Place-cells response tuning analysis**
6. **Places-cells memory analysis**

### Mouse model of antibody-mediated NMDAR Ab transfer

CSF was pooled from 10 patients with high titer IgG GluN1 Ab (all > 1:320), and 10 control subjects were used (same samples as reported in Mannara et al 2020). The presence of NMDAR Ab was verified with three different techniques, brain tissue immunohistochemistry, HEK293T cells expressing NMDAR, and cultured neurons, as reported (Ances et al., 2005; Dalmau et al., 2008). The CSF was administered for 14 consecutive days using either: i) continuous delivery via subcutaneous osmotic pumps as previously reported (Planagumá et al 2015), or ii) one daily injection for 14 days using an injector system (#C235I/Spc; PlasticsOne, Roanoke, VA) connected via a vinyl tube 10' (#C312VT-PKG; PlasticsOne, Roanoke, VA) to hamilton syringes and programmable syringe pumps (Suppl. Fig. 1). Because results were similar comparing both methods (Suppl. Fig. 2), we adopted daily injection as the new standard in all our experiments, since it decreases the risk of infections, and neck's muscles injuries generated by the subcutaneous implant of osmotic pumps. Daily injections were performed after the behavioral test to not disturb animal performance. Animals that received CSF with NMDAR Ab are further referred to as the anti-NMDAR group and those receiving CSF from control subjects as the control group.

### Calcium imaging + intraventricular infusion preparation

#### Endoscope implant

Thirty days after the adeno-associated virus (AAV) injection a second surgery was performed to implant a mini-endoscope. The surgery's goal was to implant the intracranial catheters and the mini-endoscope for calcium activity visualization in freely moving animals. The mini-endoscope is a stainless steel guide tube (1.2 mm of inside diameter x 3mm length) with a custom glass coverslip glued to one end (0.13 mm thick cover glass, Paul Marienfeld GmbH) which is the side that is in direct contact with the brain tissue. The miniendoscope holds a gradient-index (GRIN) lens (Grintech, 1mm diameter, 4mm length, n.a. 0.5). The GRIN lens is used to focus the mini-microscope used for calcium recordings. To ensure a stable attachment of the implant, small screws (18–8 S/S, Small parts) were inserted in the skull (without piercing the dura) above the cerebellum, olfactory bulbs, and somatosensory cortex to increase the implant's resistance against torque exerted by the animal while doing the task. The endoscope was then inserted at the position and angle that covered most of the flat area of the dorsal part of the hippocampal CA1 region [relative to bregma ML = 2.1(+ 77º on the coronal plane), AP = −2.2, DV = −1.1(from dura)]. To perform the implantation, a craniotomy was made using a trephine drill (~2 mm in diameter). To prevent increased intracranial pressure due to the implant, we aspirated a cylindrical volume of brain tissue equivalent to the volume occupied by the mini-endoscope. Tissue was aspirated up to the second set of fibers crossing over the CA1 area, coming from the entorhinal cortex. Each set of fibers was recognized by identifying that their orientation was ~60 degrees from the previous layer. Next, the mini-endoscope was lowered with the stereotactic manipulator centered in the coordinates displayed above and then fixed to the skull using ultraviolet-light curable glue (Loctite 4305). Metabond (Parkell) was applied around screws, the implant, and the surrounding cranium and then acrylic dental cement (Coltene, Whaledent) was placed on top of the Metabond, for the joint purpose of attaching a metal head-bar to the cranium and to further stabilize the implant.

#### Cannulae for CSF infusion preparation

In the same surgery for the endoscope, we implant bilateral cannulation (#3260PDG-1.5/SPC, Gauge 26; length 3.5mm, Plastics One, Roanoke, VA) in the two central ventricles (AP = +1.72mm; ML = +/-0.85mm; DV = -2.43mm (45º tilted to rostral direction)). These cannulas were used for intracranial, intraventricular injections of Ab. Mice recovered their normal behavior a few hours after the surgery and they returned to behavioral testing 3 days after surgery. To allow for the resolution of neuroinflammation, calcium imaging was carried out 4–5 weeks after surgery.

#### Miniscope baseplate mounting

Prior to calcium imaging recordings in our experiment, the level of GCaMP6m expression was checked daily by locating the GRIN lens within the endoscope and by approaching the mini-microscope to the focal plane of the GRIN lens in a head-fixed mouse preparation we observed the image quality.

Once global expression of calcium activity reached a steady amplitude the miniature microscope's baseplate was mounted (nVista HD, Inscopix, Inc.) by utilizing acrylic cement and the ultraviolet-light curable glue (Loctite 4305). The baseplate allows the fast attachment of the microscope to the animal for each behavioral test session. This method gives the possibility to re-attach the mini-microscope in the same focal plane to record the neuronal activity of the same neurons over the experiment length i.e. approximately three months. The baseplate includes a set of four magnets and a screw that are used for the stable attachment of the mini-microscope to the animal's head during the behavioral session.

#### Neurons' quality exclusion criteria

The distribution of firing rate within the two methods of experimentation (i.e. acute injection and osmotic pumps) was larger than across groups. We decided to apply a general algorithm to exclude outliers based on the distribution of firing rates from each of the experimental group methods (Suppl. Fig. 2). An optimization procedure was applied to a parameter set to implement our exclusion criteria. These parameters are: 1) $Fr_{max}$ , a maximum threshold on firing rates to identify and exclude fast spikers, and 2) $Score_{min}$ , a minimum threshold on cell scores to exclude low quality cells. The cost function, *J*, was defined for minimizing the difference between the mean firing rate of two animal groups which received antibodies either by osmotic or by hamilton pumps.

$$J(Score_{min}, Fr_{max}) = \sum_{days} | Fr_{control(O)} - Fr_{control(H)} |^2 + \sum_{days} | Fr_{anti\text{-}NMDA(O)} - Fr_{anti\text{-}NMDA(H)} |^2 \quad (1)$$

Here *Fr* indicates the mean firing rate of cells per session, for each group, (H) indicates the hamilton pump group, and (O) indicates the Osmotic pump group. As a result of minimizing this cost function, the maximum firing rate in CDSD was set to $Fr_{max}$ = 1.9 spikes/sec, and for NOL, $Fr_{max}$ = 2.5 spikes/sec. Moreover, the optimal Score cutoff to make the difference between groups minimal was found similar for NOL and CDSD, $Score_{min}$ = 20% of the distribution of all the scores from all cells.

#### Behavior analysis

Behavior videos were analyzed with DeepLabCut, an open-source toolbox for the estimation of the animals pose (Mathis, et al., 2018). The detected body positions were used to estimate movement trajectories as well as the animal's speed of movement (Suppl. Fig 4). According to the dominant models of hippocampal activity, the place cells' firing patterns that are displayed during exploration of an environment can be replayed during resting epochs, which are believed to play a role in the consolidation of the encoded experience; or pre-played preceding another exposure for planning future excursions in the spatial enclosure (Pfeiffer & Foster , 2013). To avoid epochs where the animals were still or without locomotion, we eliminated in the CDSD task video frames in which the animal moved slower than 1 cm/sec to avoid neuronal activity responses that do not correspond with pure spatial tuning. To evaluate cells' selectivity for spatial information, the 2D arena was divided into G=10 equal grids per dimension. The animal's location in the 2-dimensional space was computed and averaged over time bins of size $\Delta t$=250 ms. This time window was optimized to maximize spatial information. Inclusion criteria for spatial grids were met when animals visited the particular spatial bin at least four times during the whole session. Sessions in which the animal did not move faster than 1 cm/sec, for at least 5 minutes, were excluded.

#### Place-cells response tuning analyses

Similar to the process of averaging behavior trajectories over a $\Delta t$, explained above, calcium signals were also averaged over time bins of size $\Delta t$=250 ms. Then, since any neuron could exhibit varying activity patterns over different visits to the same location, rate-maps notified by r(x,y|visit), were computed to indicate the place-related activity of each individual cell, in the two-dimensional space. Here, by 'visit' we mean the visits of the animal to each spatial bin within a temporal bin used for discretization. (example schematics of rate-maps, color-coded for firing rate, are shown in figure 6). The mean rate-map is a heat-map of size G×G that is computed for each cell as the collection of its firing rate inside corresponding spatial grids and averaged over all visits of the relevant grid during the entire session.

**Identification of place-cells:** If there exists a stable relationship between a neuron's activity pattern and the animals' position in the environment, then that cell is called a place-cell (Fig. 3A). Hippocampal place-cells, on one hand, establish spatial representation in the brain, and on the other hand, can be used as readouts of the stored memories (reviewed in Moser & Paulsen, 2001). In this study, Mutual Information is used as a tool to find out how reliable a cell's activity is at different positions of the animal in the environment. In our case where the variables are the cell's spike firing rate ($v$), and the mouse 2D positions *(x,y)*, mutual information is computed as follows:

$$Mutual\ information = \sum_{x,y,v} P(x,y,v) \times log\left(\frac{P(x,y,v)}{P(x,y) \times P(v)}\right) \quad (2)$$

As stated before, $x$ and $y$ are each divided into G=10 bins so that the location probability $P(x, y)$, is a G×G matrix showing how often each spatial grid is visited by the animal. For $P(v)$, the probability of firing rates, the signal voltage range was divided into N=10 bins, and then $P(v)$ was calculated as the histogram of the signal falling in each voltage bin. $P(x, y, v)$ is the joint probability of the location and the firing rate. The measured value of mutual information for each cell was compared with the 95th percentile of a bootstrapped distribution, constructed by shuffling the firing signal relative to the animal's position in 1000 surrogate realizations. A cell that had a mutual information higher than the 95th percentile of the bootstrap distribution was identified as being significantly modulated by space, and thus labeled as a place-cell. In this work, we have defined a place-field area for a place-cell as the region of its rate-map that has values at least half of the maximum value of the rate-map. Then the mean firing rate of the rest of the rate-map is considered as the outside-field neuronal activity. Numbers (proportions) of identified place cells are displayed in Table 1.

**Spatial information analyses:** To quantify the information that a neuron carries out, I defined the Signal to Noise Ratio (SNR) measure, as:

$$SNR = \frac{var\left(\frac{1}{N}\sum_{visit}^{N} r(x,y|visit)\right)}{\frac{1}{G^2}\sum_{x,y}^{G^2} var\left(r(x,y|visit)\right)}. \quad (3)$$

The SNR is an assumption-free estimator, not dependent on any definition imposed arbitrarily, like the ones in the most commonly used descriptions of PF characteristics. The SNR is also unbiased, instead of other estimators like entropy or mutual Information (Paninski, 2003); however, it also has limitations like previously named observables because it depends on the choice of temporal and spatial discretization. We also minimized the detrimental effect of constant binning by optimizing the parameters that specify the spatial and temporal bins (see Methods, Suppl. Fig. 4).

#### Places-cells memory analysis

**Tracking cells across days:** To track cells across imaging sessions, we used an algorithm that was developed for one-photon freely-moving calcium imaging recordings, called noRMcorre (Pnevmatikakis, et al., 2017). The algorithm matches cells sorted in a given session with the cells sorted from a consecutive session. Once done for the entire experiment, it is tabulated for sessions in which each neuron was detected. The number of neurons that we can correlate over different time intervals decreases substantially the longer the interval due to a phenomenon called turn-over (Ziv et al., 2013), for which only a small fraction of neurons will reactivate on different expositions to the same environment. This fraction decays exponentially with the delay between expositions.

**Spatial memory analyses:** To assess the memory of a place-cell, we analyzed the level of similarity among the rate-maps, *r(x,y)*, exhibited at different time points. For any place-cell the more similar the rate-maps are across time, the more stable the corresponding encoded memory is. The following equation indicates how normalized cross correlation between single units of recording at time *i* and *j* were computed to evaluate similarity of rate-maps.

$$Xcorr(i,j)_{Normalized} = \frac{XCorr\,(r(x,y)_i, r(x,y)_j)}{\sqrt{max(Autocorr(r(x,y)_i))} \times \sqrt{max(Autocorr(r(x,y)_j))}} \tag{4}$$

We measured 1) the amount of correlation in the shape of place-fields, and 2) the amount of displacement at the spatial location to which a cell was maximally tuned. The cross-correlation of two rate-maps of size $G \times G$, *Xcorr(i,j)*, is itself a $(G + G - 1) \times (G + G - 1)$ matrix. The maximum value of this matrix lies on its central component (i.e. index (*G/2,G/2*)) in the case of a PC with identical fields across sessions, and deviates from the center as the PC's field shifts in space.

**Spatial memory index:** To take both the amplitude of correlation as well as the shift in the location of place-fields into account, we defined a memory index measure in the range [0 1] as: MI = Xcorr x (1 - $\text{displacement}_{\text{normalized}}$). In this equation, to normalize displacement, the amount of shift measured by Xcorr, is divided by the box size diagonal, for each experiment. It is important to notice that the MI is not an absolute quantity since higher or lower MI will depend also on the quality of the data. For this reason, we will always compare the MI with the value obtained from shuffle data which will provide us with an estimate of memory strength at the chance level, meaning, if two place fields are randomly located in the environment what is the MI at the chance that can generate. If the actual MI is above the 95% confidence interval of the shuffle data, then we are measuring actual memory with a p-value less than 0.05. Correlation in our definition of MI is Pearson's correlation of the two linearized PC's fields and the shift is the normalized shift that maximizes the overlap between the two spatial maps.

**Statistical significance of stability:** Furthermore, to generate the null hypothesis for the average place fields' stability, we used the 95% confidence interval of these two measures using a distribution constructed from 1000 pairs of random place-fields taken from a shuffled set of cells' indices. For each condition, the pair of shuffled cells were taken from the same animal group as well as the same time span that the real data came from. We considered the correlations above 95th percentile, and displacements below the 5th percentile of the bootstrap distribution to be significantly stable and different from a chance level of stability.

#### Supplementary Figures

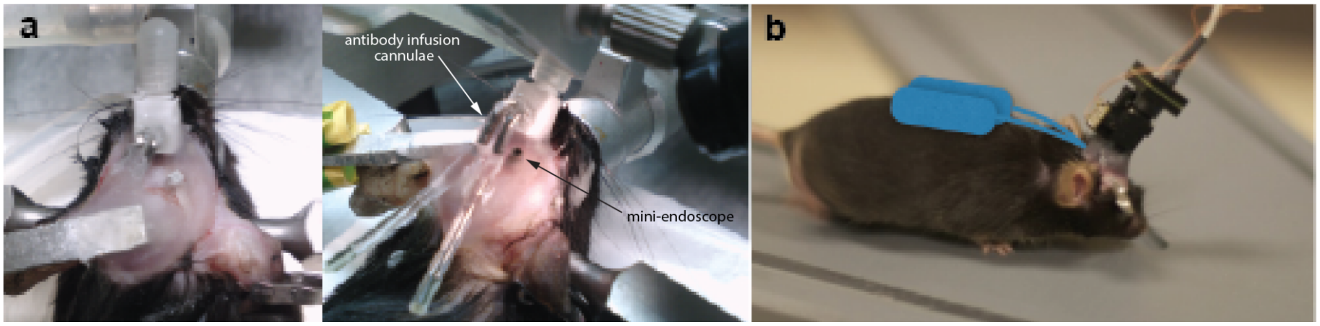


**Supplementary Figure 1: Novel preparation for calcium imaging and intracranial pharmacology.** **a)** In the same surgery we implanted a cannula for pharmacology, a mini-endoscope to later introduce the grin lens, and a head-bar to clamp the animal head during Ab infusion and mini-microscope mounting. **Right:** detail of the bi-ventricle cannula with two tubes to connect with osmotic pumps. Osmotic pumps are inserted in the back of the animal under the skin. Tubes enter the skin 3-4mm below the neck muscles, and skin is sutured to avoid infections. **b)** Picture of actual animal with mini microscope mounted + schematic of the location of the infusion pumps, to illustrate the positioning of the osmotic pumps in the back of an animal that is carrying the mini endoscope and the infusion cannula.

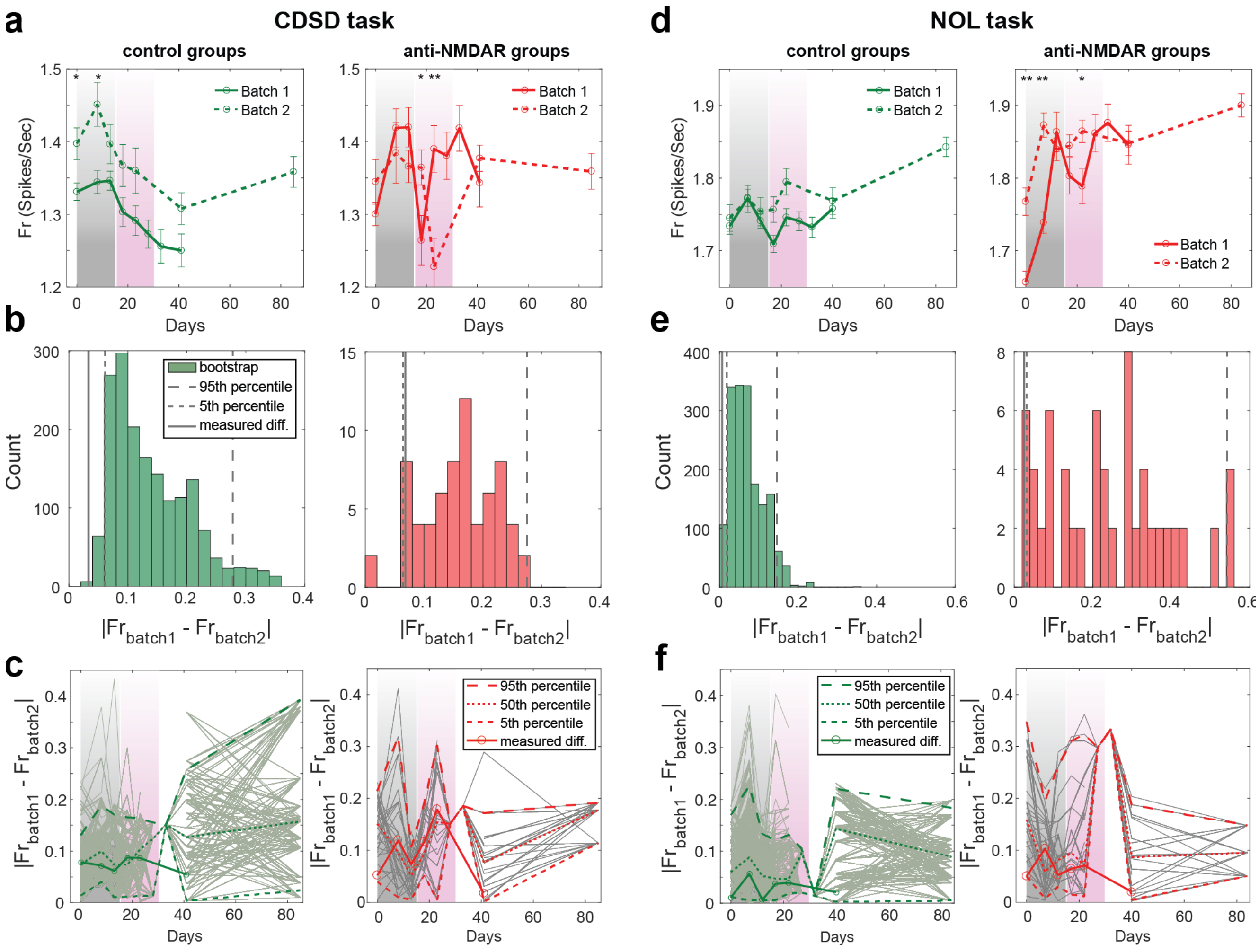


**Supplementary Figure 2: comparison between the two batches of animals using different infusion methodologies.** For CDSD task: **a)** Firing for each batch across days for control (left) and anti-NMDAR groups (right). **b)** Mean firing rate difference between the two batches for controls (left) and for the two antiNMDAR (right) groups. The distribution of mismatches is generated by shuffling the label of the animals between batches, combining all experimental days. Given the data size we cannot prove that the batches are not significantly different, but we show that they are very far from being significantly different (continuous line relative to the 5th percentile dashed line). **c)** Difference between Fr batches across days. The shuffle procedure provides a confidence interval for each experimental day. In all cases the difference between the batches is not significant, and most of the days around chance level (50th percentile dashed line). **d)**, **e)**, **f)** is the same than in a, b and c but for NOL Task.

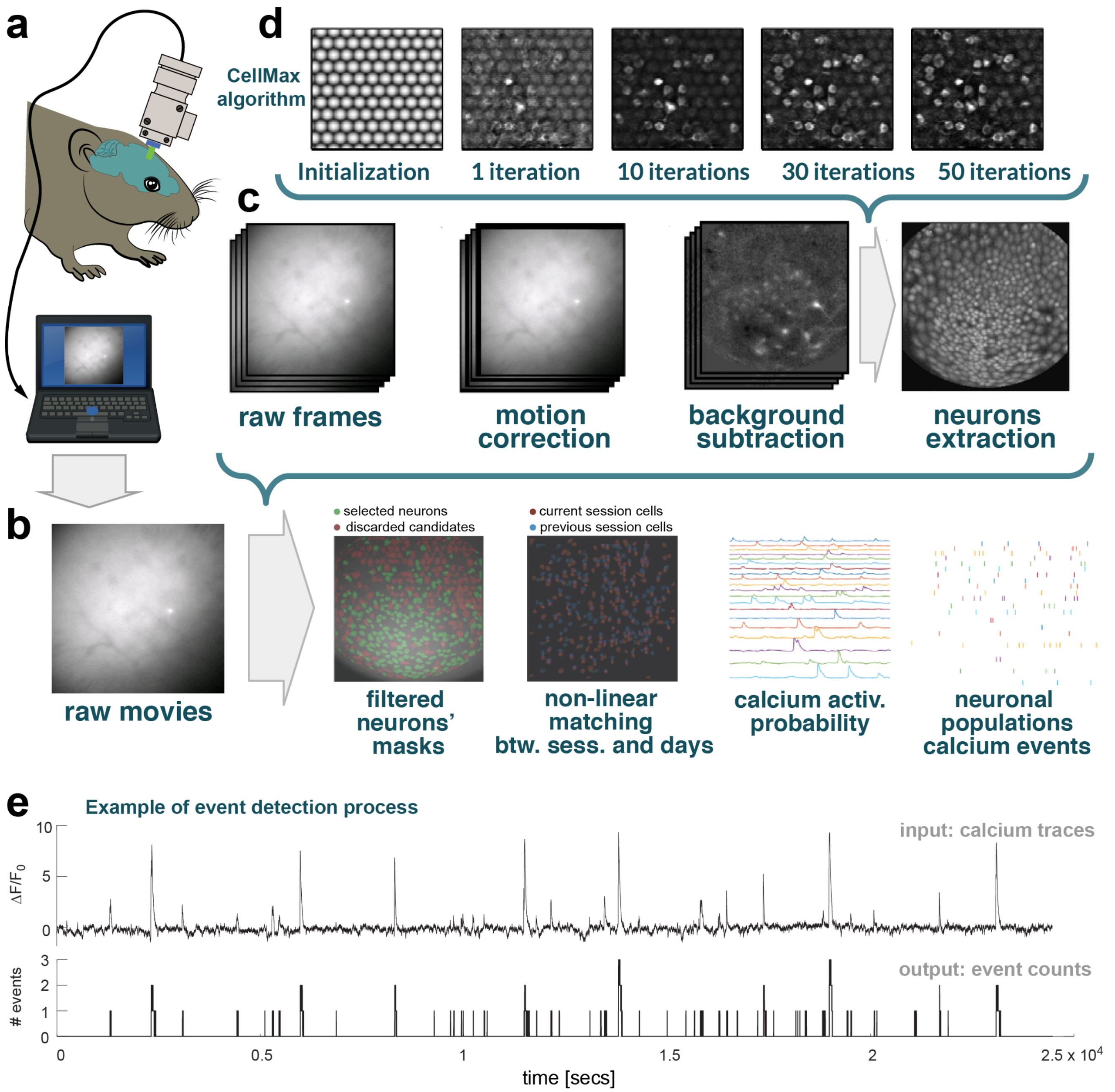


**Supplementary Figure 3**: **Neuronal activity recordings using calcium imaging methodology. a)** Schematic of the experimental process of collecting camera frames with reporting calcium activity from hundreds of neurons simultaneously. **b)** Work-flow of the implemented algorithms necessary for the processing of the calcium movies into neuron spike detection events. **c)** The first main step in the data processing is the motion correction and background subtraction to obtain a stable calcium movie of the events recorded during the different experimental behaviors (NOL and CDSD tasks). **d)** The second main processing step is to detect the shape of each individual neuron, fitting a model of photon emission to disentangle cells that overlap in the visual field (CellMax Algorithm). **e)** Once the mask of each individual neuron is obtained, then calcium traces from the average activity over each individual mask are acquired for event detection. Using the SpikeML algorithm we obtain the number of calcium spikes events per frame (50 milliseconds), which allow us to generate a firing rate

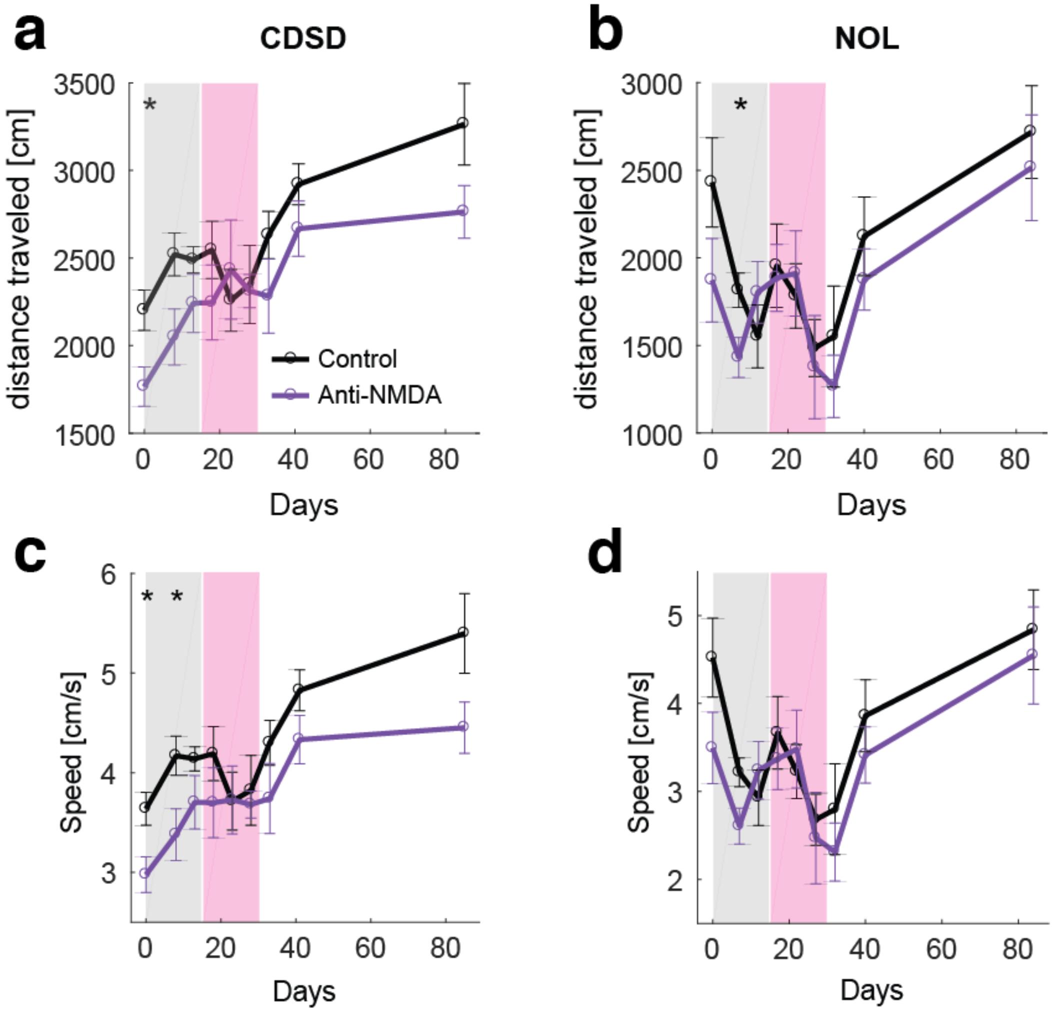


**Supplementary Figure 4: Increase in animal motivation during the experiment of ~3 months, displaying small differences between groups. a)** Distance traveled by mice performing the CDSD task increased over time, showing the willingness to engage with a task that is mostly based on foraging. Note: stars represent significant differences between groups ($p<0.01$). Two way ANOVA, p-value(time)<0.0001, p-value(treatment)=0.004, p-value(time*treatment)=0.59 (n.s.). **b)** Distance traveled by mice performing the NOL task does not change significantly over time, displaying constant motivation to solve the task Two way ANOVA p-value(time)=$10^{-4}$, p-value(treatment)=0.16 (n.s.), p-value(time*treatment)=0.59 (n.s.). **c,d)** Speed correlates with the distance traveled and also demonstrates the increase or maintenance of motivation throughout the experiment. For c) two way ANOVA, p-value(time)<$10^{-4}$, p-value(treatment)=$6 \times 10^{-4}$, p-value(time*treatment)=0.81 (n.s.). For d) two way ANOVA, p-value(time)< $10^{-4}$, p-value(treatment)=0.13 (n.s.), p-value(time*treatment)=0.67 (n.s.).

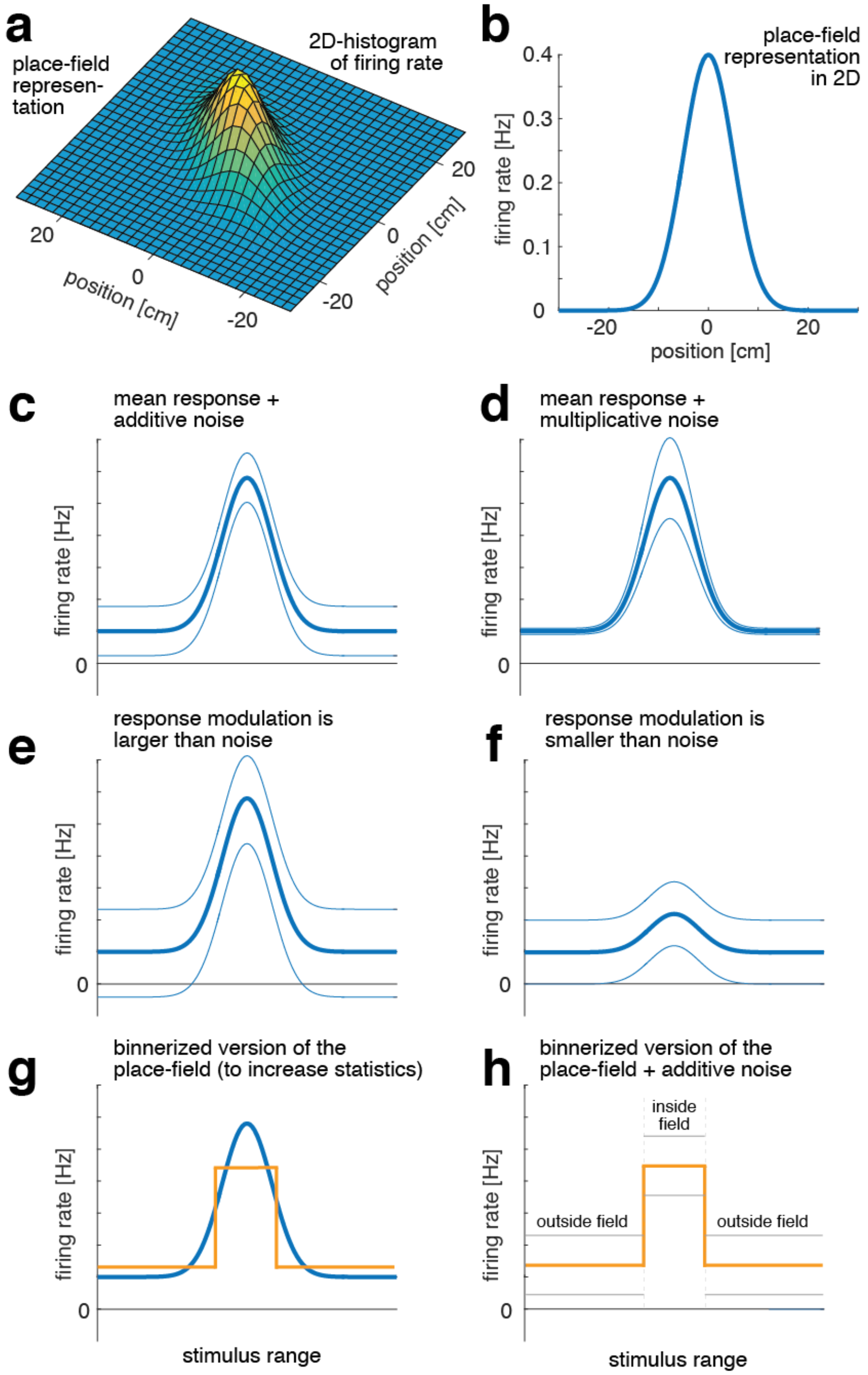


**Supplementary Figure 5: How inside/outside ratio relates to spatial code accuracy. a)** Representation of a PC's neuronal response. The firing rate is plotted for each of the spatial bins (i.e., PF). The more yellow the color is, the higher the firing rate. This is an example of one PC. Different PCs have different PF shapes and locations of the maximum firing. **b)** a 2D section of the PC representation in (a). To describe the information carried by a PC it is necessary to measure the mean response of the animal over different visits to the same spatial bin, as well as the variability of these responses. The mean response over different visits to different spatial bins is called signal (thick line). The standard deviation of the firing rate over different visits to each spatial bin is called noise (thin lines). Noise can be additive (independent of the signal) as in **c)**, or can be multiplicative (proportional to the signal) as in **d)**. For simplicity, we will use in our analysis that the noise is additive. The readout of information for each individual neuron needs to take into account, both the amplitude of the modulation of signal and noise. If the modulation of the signal is larger than the noise amplitude **e)**, the cell is spatially informative and can provide information about the animal's location. Instead, if this is the opposite case **f)**, then the neuron is not able to provide information about the position of the animal in the box. To estimate signal and noise it is necessary to measure the value of the firing rate over a large number of visits over a large number of bins. Because this is impossible to achieve with our data set, we will maximize the statistical power, by separating to main spatial regions: all the spatial bins with more than half-max of the maximum rate as the inside field, and all the spatial bins with less than half-max as the outside field. We will compute the values for the signal as the total firing rate inside and outside the field as two only values **g)**. Signal and noise are then computed for the inside and outside fields to estimate the SNR for each given neuron.

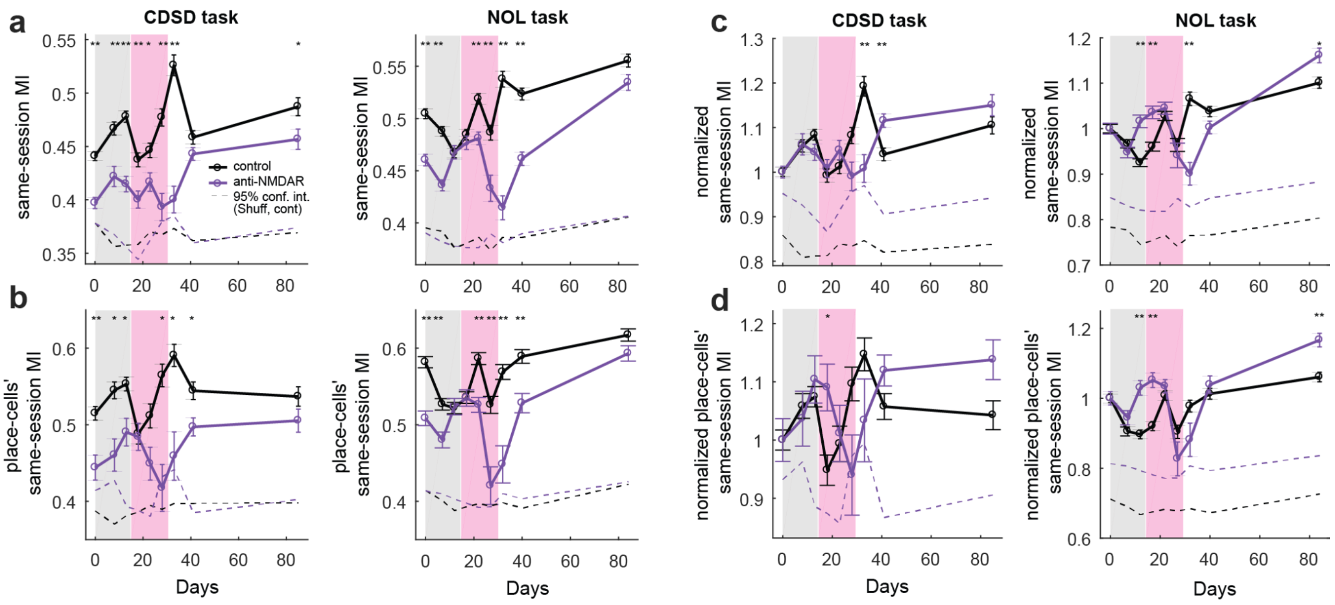


**Supplementary Figure 6: Same session memory does not change significantly between groups in a systematic manner. a)** Memory index (MI) between the first and the second half of the behavioral sessions for the two tested tasks (CDSD and NOL). We analyzed the data from the two different environments (Environment A and Environment B) in the CDSD task (memory over 10 min), and the two conditions (train and test) in the NOL task (memory over 9 min). In the case of memory on the time scale of minutes, the two maps were created using the first and second halves of the session. Memory from both sessions is highly significant relative to the 95% of the surrogate (shaffled neurons indexes) data. **b)** MI for place cells only. Due to the short duration, half sessions do not provide sufficient statistical power to observe significant changes between groups. Note: same-session MI is significantly different even on day 0 of the experiment when the antibodies have not been injected yet; for this reason we have normalized on plots c and d to reduce these differences due to small data sizes. **c)** MI for all cells, normalized relative to the amplitude on day 0. **d)** Normalized MI for place-cells only.

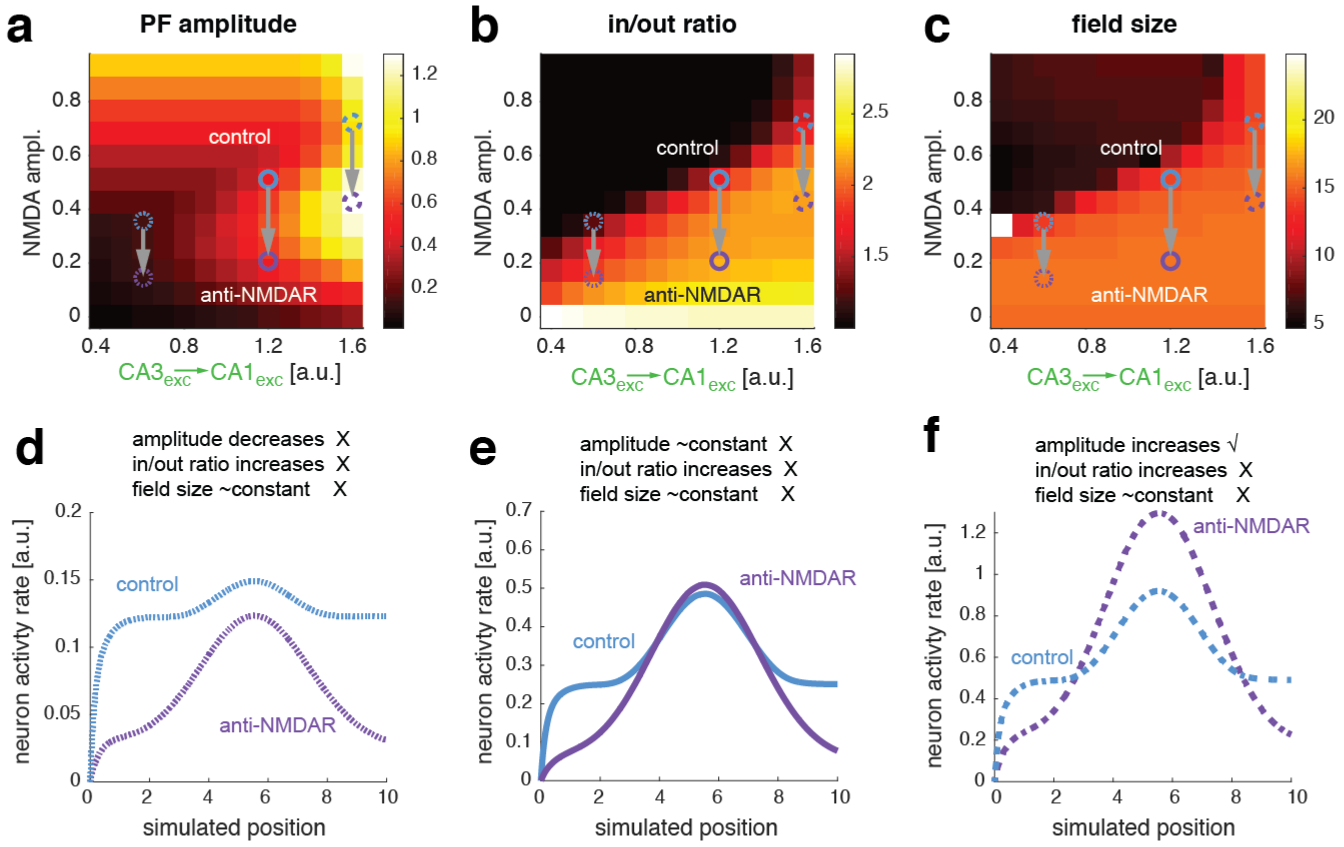


**Supplementary Figure 7: Simpler models of hippocampal circuits do not satisfy the behavior induced by the NMDAR antibodies in our data**. **a)** PF amplitude as a function of NMDAR current amplitude versus $CA3_{exc}$ to $CA1_{exc}$ synaptic strength. Each model is represented by a different set of lines (dots, solid and dashed circles) NMDAR is a constant current activated uniformly for all spatial locations. From blue to violet is the decrease in NMDAR current input due to the presence of Ab. **b)** Inside/outside ratio as a function of NMDAR current amplitude versus $CA3_{exc}$ to $CA1_{exc}$ synaptic strength. **c)** PF size as a function of NMDAR current amplitude versus $CA3_{exc}$ to $CA1_{exc}$ synaptic strength. **d)** The first model shows a decrease in PF amplitude, increase in inside/outside ratio and a maintenance of PF width, all inconsistent with our experimental observations. **e)** The second model maintains the amplitude of the PF when Ab are present, increases in the inside/outside ratio and maintains PF, all inconsistent with our experimental observations. **f)** The third model shows PF amplitude increase as in our experimental observations, but shows increase inside/outside ratio and maintenance of PF size, which are both inconsistent with our observations. Overall, none of the models that can be obtained with this type of contribution of NMDAR currents can behave as our experimental observations. NMDAR should at least depend linearly with the activity as in the model included in our manuscript (Fig. 7), to behave as in our experimental observations.

### Annex I
### Hippocampal CA3-CA1 Network Model

Pablo Jercog, Horacio G. Rotstein

November 24, 2022

#### 1 Introduction

We use a firing-rate model approach [1–3] to model the CA3-CA1 feedforward model schematically presented in Fig. 7-a. The network consists of three layers: CA3 excitatory cells (layer 1, N cells/nodes, black), CA1 inhibitory cells (layer 2, M cells/nodes, blue), and CA1 excitatory cell (layer 3, 1 cell/node, gray). For each model (i.e. set of parameters) we used a firing rate model with a (nonlinear) activation (input-output) function $F_{STNL}$, which is an adapted version of the non-saturating one presented in [Ramirez-Mahaluf et al. Cereb Cortex 2017 Jan 1;27(1):660-679] that captures the supralinear neuronal response gain at low firing rates, the sublinear response gain at higher firing rates, and is mathematically smooth (and invertible).

#### 2 Equations

The firing rate of the excitatory neurons in CA3 is denoted by $X_{1,k}$ where the sub-indices correspond to the layer $(1)$ number and the cell number $(k)$ in the layer. The firing rate for the inhibitory cells in CA1 is denoted by $X_{2,j}$ where the sub-indices correspond to the layer $(2)$ number and the cell number $(j)$ in the layer. Finally, the firing rate of the excitatory cell in CA1 is denoted by $X_3$ (layer 3).

All CA3 excitatory cells (layer 1) receive a time-dependent, Gaussian-like input and project (AMPA and NMDA) to both the CA1 inhibitory cells (black) and the CA1 excitatory cell (green). All CA1 inhibitory cells project (GABA$_A$) to the CA1 excitatory cells (blue). The output is measured in the CA1 excitatory cells. CA1 cells do not receive any external input.

The model equations are

$$\tau_{X,1} \frac{dX_{1,k}}{dt} = -X_{1,k} + F_{STNL}\left[G_{NMDA,1} X_{1,k} + F_k(t)\right] \tag{1}$$

for $k = 1, \ldots, N$

$$\tau_{X,2} \frac{dX_{2,j}}{dt} = -X_{2,j} + F_{STNL}\left[G_{NMDA,2} X_{2,j} + \sum_{k=1}^{N} J_{2j,1k} X_{1,k}\right] \tag{2}$$

for $j = 1, \ldots, M$, and

$$\tau_{X,3} \frac{dX_3}{dt} = -X_3 + F_{STNL}\left[G_{NMDA,3} X_3 + \sum_{k=1}^{N} J_{3,1k} X_{1,k} - \sum_{j=1}^{M} J_{3,2j} X_{2,j}\right] \tag{3}$$

The parameters $\tau_{X,1}$, $\tau_{X,2}$ and $\tau_{X,3}$ are the time constants. The parameters $G_{NMDA,1}$, $G_{NMDA,2}$ and $G_{NMDA,3}$ capture the effects of NMDA. The parameters $J$ denote the connectivity. We adopt the convention that the first sub-index indicates the layer and cell number of the postsynaptic node and the second sub-index indicates the layer and cell number of the presynaptic node (e.g., $J_{2j,1k}$ is the connectivity from cell $k$ in layer $1$ to cell $j$ in layer $2$). We used the following parameter values: $\tau_{X,1} = \tau_{X,3} = 20$, $\tau_{X,2} = 10$, $G_{NMDA,1} = G_{NMDA,2} = G_{NMDA,3} = 0.9$, $J_{2j,1k} = 0.2/N \; \forall \; j,k$, $J_{3,1k} = 1.2exp(-(k-N/2)/5^2)$, $J_{3,2j} = 0.8/M \; \forall \; k$

The input-output function $F_{STNL}$ (smooth threshold nonlinear) is defined as

$$F_{STNL}(x) = \begin{cases} \eta/[1+exp(-(x-x_{th})/x_{slp})], & \text{if} \quad x < 0, \\ x^2+\epsilon, & \text{if} \quad 0 \le x \le 1, \\ 2\sqrt{x-3/4}+\epsilon, & \text{if} \quad 1 < x. \end{cases} \tag{4}$$

We used $x_{th} = 0, x_{slp} = 1, \eta = 0.0001$, and $\epsilon = 0.5\,\eta$.

The function $F_{STNL}$ is a smooth extension of the threshold nonlinear function used in [4] and a smooth nonlinear extension of the threshold-linear input-output functions used in rate models (see [3] for details).
